# An atlas of shark developing telencephalon reveals ancient origin of basal progenitors and Cajal-Retzius cells

**DOI:** 10.1101/2025.05.08.652169

**Authors:** Idoia Quintana-Urzainqui, Tobias Gerber, Phillip A. Oel, Leslie Pan, Nikolaos Papadopoulos, Z. Gülce Serka, Ana Verbanac, Maite Börsig, Dorinda Torres-Sabino, Isabel Clara Rollán-Delgado, Luca Santangeli, Henrik Kaessmann, Detlev Arendt

## Abstract

The emergence of predation and associated complex behaviors in jawed vertebrates (gnathostomes) were major driving forces in brain evolution. To gain insight into the neuronal complexity of the last common gnathostome ancestor, we studied the development of the shark telencephalon by combining single-cell and spatial transcriptomics. Our findings suggest an ancient origin of pallial neurogenesis, including basal progenitors, which were previously only identified in tetrapods. We find evidence for migrating gabaergic neurons populating the dorsal pallium, where we observe incipient layering. Finally, we provide evidence for the existence of Cajal-Retzius cells in the developing shark telencephalon, with a conserved molecular profile and tissue localization relative to mammals. Our analyses revealed that major features of the vertebrate pallium arose much earlier than previously thought, in the gnathostome lineage.

## Introduction

The emergence of jawed vertebrates (gnathostomes), around 450 million years ago, was a major step in vertebrate evolution. The appearance of new ecological niches exerted profound changes in body and brain organization and complexity, likely enabling higher processing speed and improved associative learning. However, the exact path of how and when the telencephalon expanded and complexified during evolution remains unsolved, due to the lack of comparative data from basal vertebrate branches. Actinopterygians (ray-finned fishes) and, in particular teleost fish, experienced additional rounds of genome duplication and exhibit unique patterns of brain development and morphology, which complicate direct comparisons and evolutionary inferences. On the contrary, cartilaginous fish are slow evolvers and show conserved features in brain development (*1–4*), making them excellent models to investigate conserved traits and pinpoint ancient innovations that have shaped early gnathostome brains.

Developmental comparisons are particularly well-suited for reconstructing evolutionary history because evolutionary change is often driven by changes in development. Furthermore, embryonic processes are more likely to retain ancestral features that are modified or even absent in adults. Therefore, the study of development is central to our understanding of how neuronal diversity and complexity evolved in vertebrate brains. In this study, we investigate the cellular landscape of the shark telencephalon to address the developmental origins of the gnathostome brain. The telencephalon as a key centre of sensory integration and associative learning is highly relevant to understanding vertebrate brain evolution; yet its step-wise elaboration and expansion is poorly understood. Our focus is on key aspects of telencephalic development that appear linked to these changes.

While the neuron types derived from the ventral side of the telencephalon (subpallium), appear largely conserved, the structure and composition of the dorsal side (pallium) has proven highly variable, which has so far precluded insights into its evolutionary path (*5–9*). A longstanding question concerns the true evolutionary origin of specialized neural cell types such as Cajal-Retzius (CR) cells. CR cells are critical for cortical development in mammals, where their key function is to regulate laminar organization through reelin signaling. While CR cells have been extensively studied in mammals, their evolutionary history remains unclear.

In this study, we investigate the developmental landscape of the shark telencephalon to address fundamental questions about telencephalic evolution. Using transcriptomic profiling, trajectory analysis and tissue mapping during embryonic development, we show that the shark embryonic telencephalon shows a degree of diversification of territories comparable to that of tetrapods, including an emerging dorsal pallial area receiving gabaergic migrating neurons and containing a cell type homologous to pallial amplifying basal progenitors. Most strikingly, we find Cajal-Retzius-like cells spatially correlating with some degree of a layered organization. Our results indicate that features previously considered to be mammalian or amniote innovations were already present in gnathostome ancestors and most likely contributed to the complexification of the telencephalon coinciding with the sophistication of their brains and behaviours.

## Results

### Single-nucleus transcriptomics captures shark telencephalic developmental trajectories

To construct a single-nucleus transcriptomic dataset of the developing shark telencephalon, we microdissected the telencephalon (including the olfactory bulbs) from three catshark embryos (*Scyliorhinus canicula*) at stages 29,30 and 31 (10) representing stages of neurogenesis and neuronal differentiation (1) (Fig. 1A) (equivalent to mouse E12-14). We isolated and dissociated the nuclei and ran three independent captures using 10X Chromium Next GEM Single Cell 3’ v3.1, which yielded 21,686 high-quality nuclear transcriptomes after bioinformatic quality control filtering with Seurat (11) and integration using Harmony (fig. S1) (12). Most cells expressed neuronal markers, with smaller non-neuronal populations identified as astrocytes (gfap, s100, sox9), choroid plexus cells (rbm47, glis3), and vascular-associated cells (flt1, piezo2) (Fig. 1B, fig. S1).

**Figure 1.**
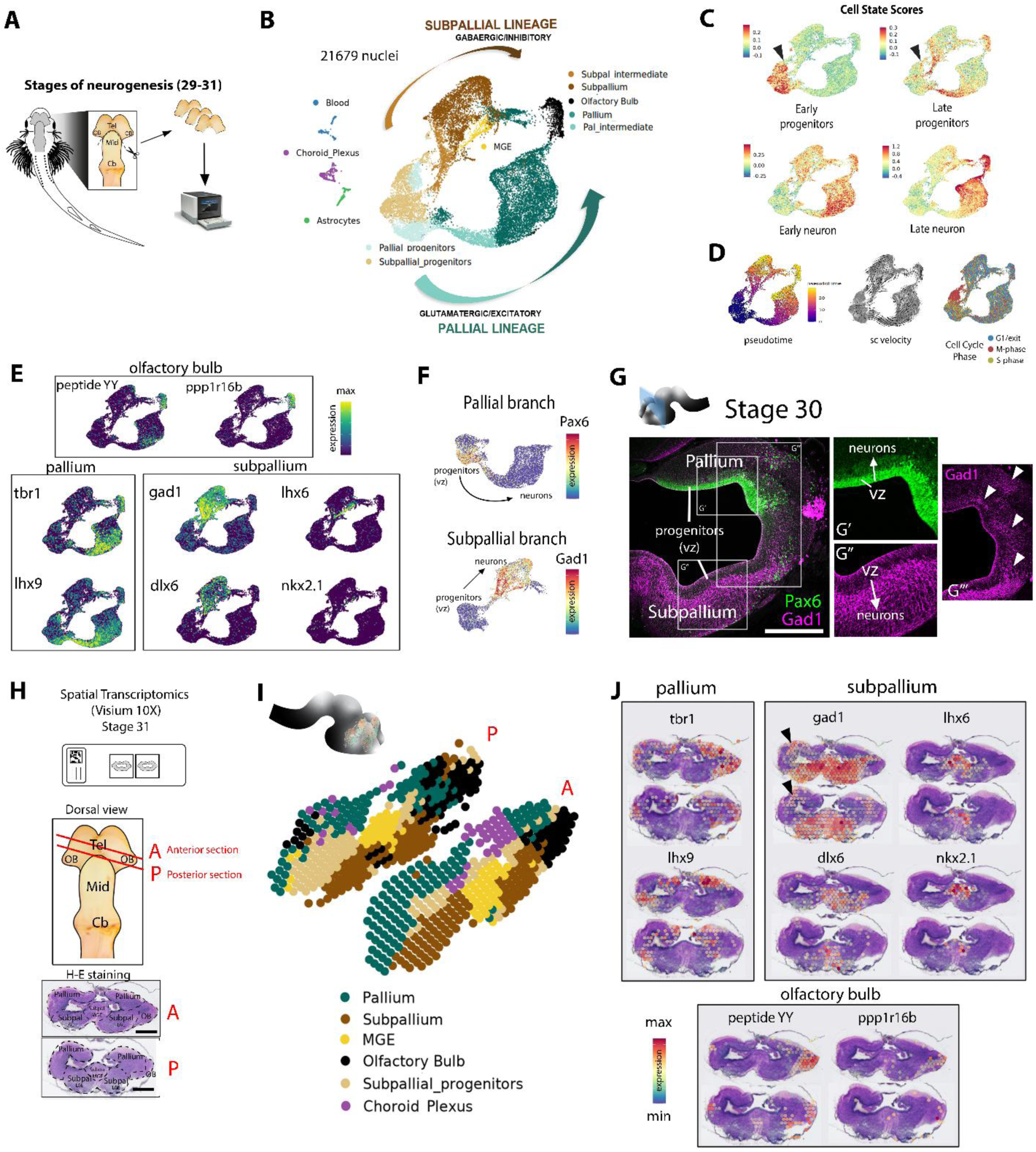
Cell type atlas of the developing shark telencephalon. (**A**) Schema of the telencephalic dissections from three independent embryos at stages of neurogenesis 29, 30 and 31 (according to (*10*)). (**B**) Umap representation of the single nucleus dataset showing the main cell types and lineages identified. (**C**) Cell state score analysis reveals the presence of early and late progenitors, early and late neurons depicting an axis of differentiation from left to right. (**D**) Pseudotime, Velocity and Cell Cycle score analysis confirm the presence of a developmental trajectory progression in our data. (**E**) Featureplots showing the expression of pallial, subpallial and olfactory bulb enriched genes. (**F**) Featureplots showing the expression of pax6 in the pallial branch and gad1 in the subpallial branch matching that of the one showed in (G). (**G**) Fluorescent immunohistochemistry at stage 30 in a transverse section showing the expression of Pax6 and Gad1 antibodies. (**G’-G’’**) Details of the pallium where Pax6 is expressed at high levels in the vz and gradually decreases its expression as neurons differentiate away from the ventricular surface; Gad1 is not expressed at the vz and strongly expressed in maturing subpallial neurons. (**H**) Schema of the spatial transcriptomics experiment (Visium 10x) showing the level of the transverse sections. (**I**) Representation of the two Visium 10x slides showing the spatial prediction of the main clusters shown in (B). (**J**) Spatial expression of the same genes showed in (E ). Arrowheads indicate gabaergic signal located at pallial areas. Abbreviations: Cb, cerebellum; Mid, midbrain; MGE, medial ganglionic eminence; Tel, telencephalon; OB, olfactory bulb; vz, ventricular zone. Scale bar: 200 μm in (G), 300 μm in (H).

To investigate whether our data contained neurons in progressively different developmental stages, we calculated cell-state scores using gene sets diagnostic of telencephalic cell states in mice (*13*). Despite the large evolutionary distance between shark and mice, this approach revealed a clear differentiation axis, ranging from early progenitors to maturing neurons (Fig. 1C). This was further supported by pseudotime analysis (Monocle3(*14*)), RNA velocity (*15*), and cell cycle scoring (Fig. 1D), collectively indicating that gene expression patterns along telencephalic trajectories are conserved between sharks and mouse.

The vertebrate telencephalon comprises two major areas: the dorsal telencephalon or pallium (origin of excitatory glutamatergic neurons) and the ventral telencephalon or subpallium (origin of inhibitory gabaergic neurons). Our dataset captured two distinct developmental branches corresponding to pallial and subpallial differentiation, emerging from their respective early progenitor clusters (Fig. 1B). Pallial markers were restricted to the lower branch in the umap, while subpallial markers defined the upper one (Fig. 1E). We found early pallial progenitors express pax6, which is gradually downregulated as progenitors differentiate into pallial neurons (curved arrow in Fig.1F). Gad1 (GABA-producing enzyme) transcript is not expressed in early subpallial progenitors and sharply upregulates as they differentiate into subpallial neurons (arrow in Fig.1F). The same expression pattern trend was observed in transverse sections of stage 30 catshark telencephalon by fluorescent immunohistochemistry (Fig. 1G), where early progenitors are located in the ventricular zone (vz) and, as neurons differentiate they migrate away from the vz, towards subventricular positions (arrows in Fig.1G’,G’’).

Guided by the expression of pallial and subpallial markers we annotated 11 clusters, representing broad cell types or cell states (Fig. 1B). Cluster markers for this resolution are included in data S1. We further subdivided the dataset at higher resolution and identified 33 candidate cell types (Fig.S1, dataS2).

To spatially map gene expression during shark telencephalic development we generated a spatial transcriptomic dataset on stage-31 telencephalic transverse sections using the Visium 10X platform (Fig. 1H). The sectioning plane was slightly oblique relative to the mediolateral axis (red lines in Fig.1 H), resulting in the right hemisphere displaying more caudal structures, such as the olfactory bulb, while the left hemisphere in the same section showing more rostral regions. We predicted the most likely cluster identity for each spot in the Visium 10X array, based on the main transcriptional profiles identified from the single nucleus data as reference signatures (Fig. 1I, fig. S1), using CIBERSORTx (*16*). The overall prediction agreed with general pallial, subpallial and olfactory bulb markers, which showed complementary dorsal, ventral and lateral distribution (compare Fig. 1E with I,J). Additionally, gabaergic signal was detected within pallial territory (arrowheads in Fig. 1J), likely representing migratory subpallial-derived cells, as observed at earlier stages (Fig. 1G’’’), and previously reported by us (*1*).The medial ganglionic eminence (MGE), a pan-vertebrate-conserved subpallial structure, which develops into the pallidal component of the basal ganglia, was localized to a ventromedial bulge in the subpallium (Fig. 1I), as indicated by the prediction and distribution of MGE-specific gene markers lhx6 and nkx2.1 (Figs. 1I,J). Interestingly, the shark MGE appears as a single medial bulge rather than bilateral structures seen in other jawed vertebrates.

Overall, our single-nucleus and spatial transcriptomic datasets provide a comprehensive survey of shark telencephalic development, recapitulating pallial and subpallial lineage progression and identifying major differentiated neuron types (Fig. 1B).

### Trajectory study of the catshark subpallium reveals three main differentiation pathways for gabaergic inhibitory neurons

To investigate the shark developing inhibitory cell types and their associated developmental trajectories, we bioinformatically isolated and reclustered cells from the subpallial branch. Three major branches emerged from the progenitor’s cluster, confirmed by trajectory inference algorithms (Monocle 3) (Fig.2A). We predicted gene expression dynamics along each subtrajectory by identifying subpallial clusters and their corresponding differentially expressed genes (methods) (Fig. 2B). Neuronal identities of each branch were assigned by integrating trajectory inference with the spatial distribution of predicted clusters (Fig.2C) and their enriched gene signatures (Fig. 2D). We observed that early gabaergic neuron differentiation proceeds through an intermediate state, marked by the upregulation of transcription factors Dlx1 and Isl1 (Fig. 2B, fig. S2). From this state, the three distinct branches emerge. The first one was predicted to give rise to interneurons in the medial bulge of the subpallium, which corresponded to the MGE/subpaliall septum trajectory based on anatomical position and gene expression (Fig. 2B,C). This branch was characterized by the specific expression of Lhx6 (MGE part) and Nxph1 (septal part) (Fig.2D).

**Figure 2.**
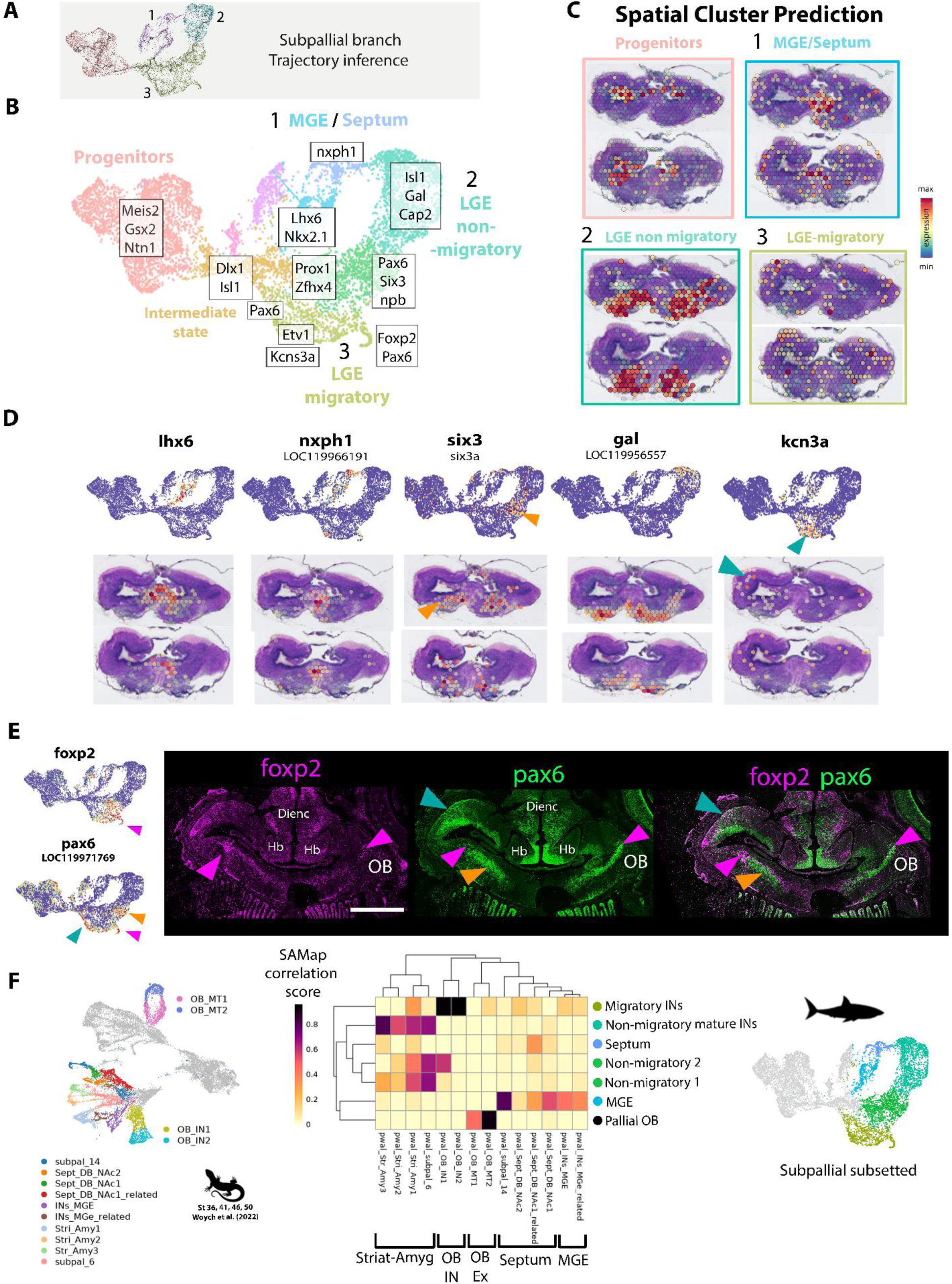
Single nucleus combined with spatial transcriptomics revealed subpallial trajectories, neuron subtypes and their anatomical positions in the shark developing telencephalon. (**A**) Subsetting of the subpallial neurons and reclustering revealed three main branches or trajectories predicted with Monocle3. (**B**) Schema showing the three subpallial trajectories and the main genes upregulated along them. (**C**) Spatial prediction of the main clusters projected in the Visium dataset. (**D**) Expression of selected genes in the umap (upper part) and in the Visium slides (bottom). (**E**) Expression of foxp2 and pax6 in the umap (left) and in the tissue shown by Hybridization Chain Reaction (right). Blue arrows point to migratory interneurons in pallial areas. Pink arrows point to dorsalLGE neurons potentially migrating towards the olfactory bulbs. Yellow arrows point to LGE-neurons that do not migrate tangentially, but stay in the subpallial territory. (**F**) Cross-species comparison with SAMap between shark and salamander developing telencephalic datasets. Only clusters colored in the umaps were included in the comparison. Abbreviations: Amy, amygdala; DB, diagonal band; Dienc, diencephalon; dLGE, dorsal lateral ganglionic eminence; Hb, habenula; IN, interneurons; LGE, lateral ganglionic eminence; MGE, medial ganglionic eminence; Nac, nucleus accumbens; OB, olfactory bulb, Sept, septum; Str, Striatum. Scale bar: 300 μm in (E).

Neurons belonging to the second subpallial branch were predicted to be distributed primarily within the subpallium, with no overlap observed with the MGE area (Fig. 2C). These cells correspond to LGE-derived interneurons that remain within the subpallium and do not undergo tangential migration to other regions. Their developmental trajectory is defined by sequential upregulation of Prox1, Zfhx4, Six3 and Pax6 (yellow arrows, Fig. 2D,E), culminating in populations expressing Cap2, Galanin and Islet1 (Fig.2D, fig.S2 Spatially, these genes also show sequential expression across the subpalliall wall (Fig.2D, fig. S2), probably reflecting radially migrating interneurons that populate a U-shaped anatomical area called the area superficialis basalis (ASB), a conspicuous mass of cells in the subpallium of all chondrichthyans, whose identity and homology to other vertebrates remain to be fully elucidated. (*17–19*).

Subpallial neurons within the third branch were not restricted to the subpallial territory but predicted to be located across different telencephalic areas, including the pallium (Fig.2C). We interpreted this distribution as indicative of tangentially migratory interneurons. Along this branch, neurons specifically upregulate etv1, kcns3 and Pax6 (Fig.2B; blue arrowheads; Fig.2D,E; fig.S2), Foxp2 and Pax6 co-expressed predominantly at the tip of the branch (purple arrowheads, Fig.2E). The spatial expression of these markers suggests the presence of two principal interneuron subpopulations: 1) Pax6 and Kcn3-positive but Foxp2-negative cells, corresponding to interneurons migrating into pallial regions (blue arrowheads, Fig.2D,E), like previously described in Fig. 1G,J (arrowheads) and in (*1*); and 2) Foxp2 and Pax6 co-expressing interneurons. To determine the precise spatial location of the latter we performed HCR for these two markers and found that this population probably corresponds to dorsal LGE-derived interneurons migrating towards the olfactory bulb (purple arrowheadS in Fig.2E).

Finally, we performed cross species comparisons using SAMap (*20*). We confronted our shark subpallial clusters with a dataset from the salamander developing telencephalon (*5*), in which main subpallial neuronal types were annotated. As a quality control, we included in the comparison pallial olfactory bulb clusters from both species, which highly correlated with each other and with no other clusters (Fig.2F). The shark MGE cluster highly correlated with MGE developmental trajectory in the salamander, as well as septal areas. Shark non-migratory clusters, both developing and mature parts, showed high correlation with striatal and amygdalar clusters of the salamander. Our results support the interpretation that the U-shaped subpallial structure mentioned above (the BSA) comprises the striatum and specific regions of the subpallial amygdala in Chondrichthyans which is a long-debated question in shark neuroanatomy (*3*, *17–19*, *21–23*).

In contrast, the shark migratory cluster found overall high correlation with salamander olfactory bulb interneurons. Judging by their position in the tissue, it is likely that a subset of the shark migratory interneurons (those co-expressing Pax6 and Foxp2) correspond indeed with migratory neurons towards the olfactory bulb derived from the dorsal LGE (dLGE) (pink arrows in Fig.2E). We hypothesize that the shark migratory cluster probably contains a second migratory population directed towards the pallium (Pax6-positive, Foxp2 negative), which did not find any clear correlation in the salamander dataset.

In summary, we have found three main developmental inhibitory neuron trajectories, likely conserved across vertebrates: 1) a MGE trajectory and its closely related subpallial septum interneuron type; 2) LGE-derived striatal and amygdalar neurons and 3) dLGE-derived olfactory bulb neurons. Additionally, we observed a distinct migratory interneuron type targeting pallial areas, which exhibits divergent gene expression profiles. Notably, this pallial-bound migratory interneuron population in sharks appears to originate from the LGE rather than the MGE, contrasting with the mammalian paradigm where pallial interneurons primarily arise from the MGE. Moreover, pallial-bound migrating interneurons do not express Pax6 in mammals, suggesting this population might be shark/cartilaginous fish-specific.

### Study of pallial developmental diversity uncovers four main excitatory neuron identities, and identifies a potential dorsal pallial region homolog

The vertebrate pallium is one of the regions of the brain that have experienced the highest variability across animal groups and one of the most challenging for cross-species comparisons. The functional roles and homologies of the pallial subdivisions in cartilaginous fish remain largely unknown, particularly regarding their correspondence to pallial regions in other vertebrates. To shed light into the shark pallial cell identities and anatomy, we focused on the main pallial branch. We excluded from this analysis the pallial olfactory bulb and two small subpopulations that cluster closer to the subpallium, which will be examined separately.

In contrast to the subpallial lineage, the pallial trajectory did not show distinct branching but instead formed a more continuous landscape of cell states. Trajectory inference with the Monocle3 algorithm predicted two main paths diverging from early pallial neurons and converging at their endpoints (Fig. 3A). We observed that genes enriched on each pallial trajectory in the umap distributed into two different anatomical areas corresponding to the ventrolateral pallium (nr2f1, epha7) and the dorsal-medial pallium (sall1, satb1) (Figure. 3A,C). Further subclustering showed the existence of at least four robust clusters representing four molecularly distinct cell types (Fig. 3D). Spatial score predictions for these clusters showed that, while there was some anatomical overlap, the distribution of their high-score areas ordered in a progression from more medial to more ventral regions, which prompted us to define them as medial, dorsal, lateral and ventral pallium (MP, DP, LP, VP), respectively (Fig.3E-G). The distribution of top enriched genes for each of these clusters confirmed their identity in the umap and position in the tissue (Fig.3H, fig.S3A). After closer examination of these cluster markers distribution we observed that, at rostral levels, all four pallial areas are evident, whereas at more caudal levels—specifically near the olfactory bulb—the medial pallium predominates, with only remnants of the dorsal pallium detectable (Fig. 3F). This refined regionalization provides a valuable reference for interpreting the neuroanatomy of the shark pallium.

**Figure 3.**
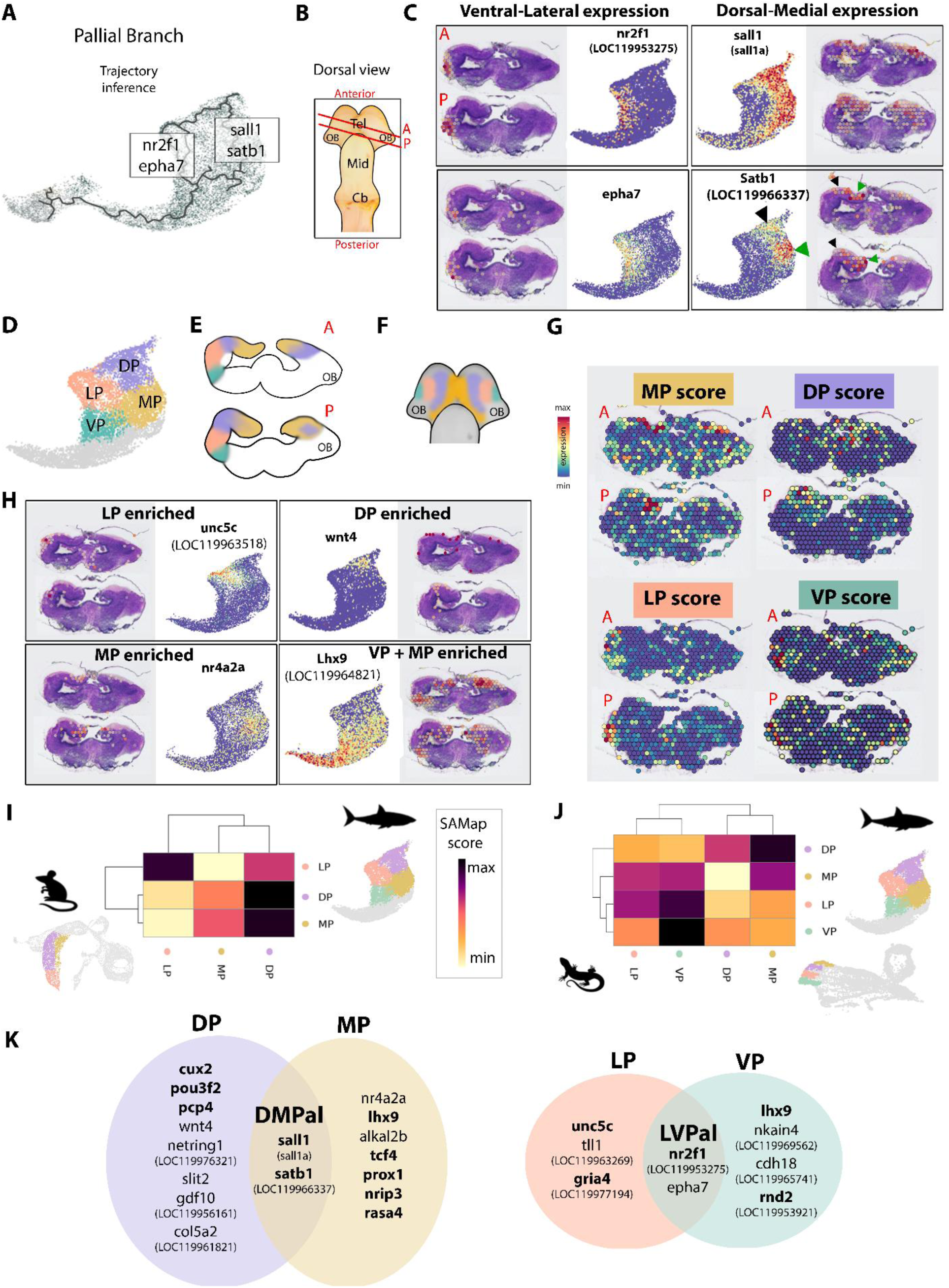
Study of the pallial trajectory and pallial subtypes. (**A**) Schema showing the subsetted pallial branch and trajectory inference with Monocle3. (**B**) Schematic representation of a dorsal view of the shark embryonic brain showing with red lines the level of the two transverse sections processed for spatial transcriptomics. (**C**) Expression pattern of the genes showed in (A), in the Visium 10x sections and in the pallial branch of the Umap. (**D-F**) Schematic drawing showing clustering of the pallial mature neurons into four main cell identities and their anatomical localization in a transverse sections (E) and dorsal telencephalic view (F). (**G**) Spatial score predictions for the four pallial areas shown in (D). (**H**) Featureplots and spatial distribution for selected genes that showed spatially-restricted expression into the different pallial areas. (**I**) Cross-species comparison across shark and mouse pallial cell types. (J) Cross-species comparison across shark and salamander pallial cell types. (**K**) List of genes enriched on each of the pallial areas defined in shark. Genes in bold indicate their expression in the equivalent pallial region of the mouse. Abbreviations: DMP, dorsalmedial pallium; DP, dorsal pallium; Mid, midbrain; LP, lateral pallium; LVP, lateralventral pallium; MP, medial pallium; OB, olfactory bulb; Tel, telencephalon; VP, ventral pallium. Red letters “A” “P” indicate “anterior” and “posterior”.

We next aimed to compare the pallial territories identified in shark with pallial maturing neurons characterized in other vertebrates. To this end, we compared our dataset with those from mouse and salamander developing pallium at similar stages. For the salamander, we utilized the pallial annotations as provided in the original study (*5*). For the mouse, we annotated ventral, lateral, medial and dorsal pallial regions based on well-established marker genes (fig. S3C). We imported two mouse datasets: a telencephalic dataset from embryonic day 12.5 (E12.5) (*13*), and pallial dataset from E13.5-14.5 (*24*), and integrated them (fig. S3B). We observed minimal correlation between maturing shark pallial cell types and E12 mouse pallial cell types, whereas the correspondence was substantially greater with the E13.5–14.5 mouse dataset (fig. S3D). We therefore subsequently performed the comparison between shark pallial types and mouse E13-14 pallial neurons. This mouse dataset was derived from an Emx1-Cre sorted line (*24*), which selectively labels pallial excitatory neurons while excluding ventral pallial populations; accordingly, we excluded the shark ventral pallial cluster from the cross-species comparison with mouse. When confronting SAMap correlation scores between shark and mouse mature pallial clusters, we observed a strong correspondence specifically between the lateral pallial cell types of both species (Fig. 3I). Both the dorsal and medial pallial clusters in shark showed the highest correlation with the mouse dorsal pallium, moderate correlation with the mouse medial pallium, and little to no correlation with the mouse lateral pallium. Overall, these results indicate that the embryonic shark pallium contains correlates of both dorsal and lateral pallial regions, likely homologous to those in mouse.

We next applied the same comparative approach to pallial clusters between shark and salamander (Fig. 3J). With the exception of the shark medial pallium, which showed similarity to medial, ventral, and lateral salamander clusters, the other pallial regions displayed more distinct matching patterns. Notably, there was a strong mutual correlation between shark and salamander ventral pallial clusters. The shark lateral pallium correlated most strongly with both lateral and ventral salamander clusters, whereas the shark dorsal pallium showed correlations with dorsal and medial pallial sectors of the salamander. Overall, we identified a clear trend in which dorsal–medial pallial sectors correlated with one another across species, and latero–ventral sectors showed a similar alignment. These patterns suggest conserved field-level homologies that may reflect ancient developmental programs guiding the differentiation of equivalent pallial areas.

We next characterized molecular signatures of shark pallial regions (Fig. 3K) by identifying differentially expressed genes for each region (0.5 logfold change) (Fig. 3K, dataS3). Genes in bold indicate shared expression with mouse (dataS4), suggesting ancestral transcriptional signatures. Notably, the shark dorsal pallium is enriched for Cux2 and Pou3f2, key regulators of upper-layer cortical neurons in mammals (*25*).

Overall, our results indicate that the shark dorsal pallium shares correlations and gene expression features with the dorsal and medial pallium of both mouse and salamander, suggesting that this field was already present in the last common ancestor of jawed vertebrates. Our findings provide a molecular an anatomical framework of the excitatory neuron types in the shark pallium, paving the way to unravel chondrichthyan forebrain organization. More broadly, they support the early emergence of a distinct dorsal pallial field in vertebrate evolution and raise further questions about additional features that may have originated at this time.

### Studying progenitor diversity reveals the presence of basal progenitors in the shark pallium

The dorsal pallium has been the focus of increased interest among neurobiologists as its evolutionary enlargement gives rise to the expanded mammalian neocortex. One of the main mechanisms known to give rise to this expansion during development is the emergence of indirect neurogenesis: the interplay between apical (AP) and basal progenitors (BP) (*26*). In mammals, APs divide at the apical surface of the ventricular zone (vz) (Fig.4A), while BPs, which arise from asymmetric divisions of APs, migrate to the subventricular zone (svz) for mitosis (Fig. 4B), where they exponentially increase the number of neurons produced by a single AP. Cell-state score analysis predicted two broad progenitor classes in the developing shark telencephalon: early progenitors and late progenitors (Fig. 1C), which correspond to mammalian APs and BPs, respectively. The potential presence of BPs in the pallium of sharks has significant evolutionary implications, as this would indicate that the emergence of pallial indirect neurogenesis preceded the expanded mammalian neocortex. Genes associated with early APs, such as gfap and notch family members, were enriched in clusters predicted to be APs (Fig. 4C). Within this population, we identified pallial APs expressing pax6 and lrp4, and subpallial APs enriched for meis2a and ntn1 (Fig. 4C, fig.S4A), consistent with patterns observed in other vertebrates(*13*).

**Figure 4.**
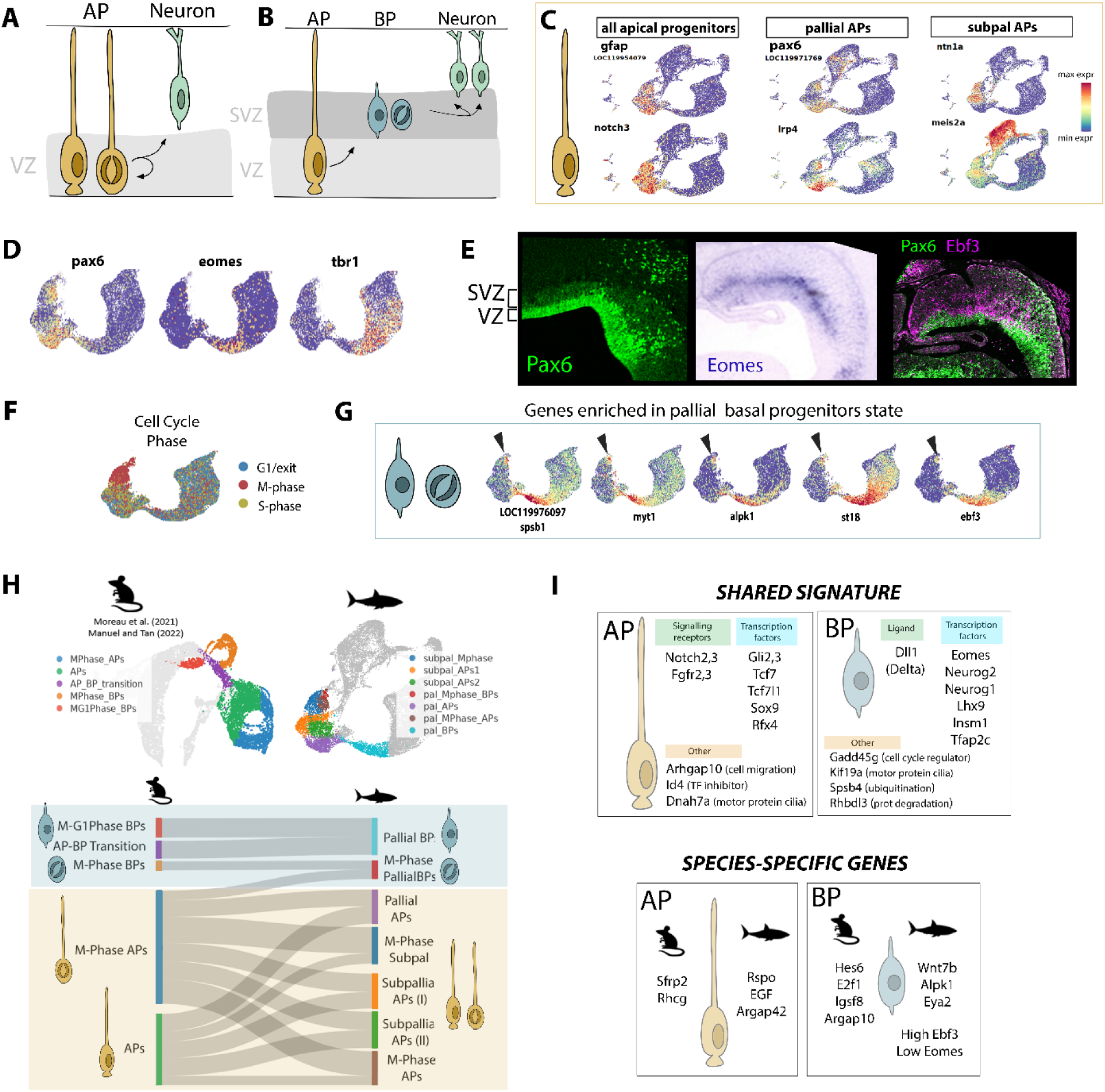
Profiling progenitor cell states in the catshark telencephalon. (A-B) Schematic drawing of direct neurogenesis (A) and indirect neurogenesis through basal progenitors (B). (**C**) Feature plots showing the expression of markers of the main types of apical progenitors. (**D**) Feature plots showing the sequential expression of pax6, eomes and tbr1 along the early pallial differentiation trajectory. (**E**) Tissue expression of Pax6 protein (left), eomes transcript (middle), pax6 and ebf3 transcripts (right). (**F**) Feature plot showing cell cycle phase scoring in the progenitors and pallial branch. Cells undergoing mitosis cluster together (red cluster). (**G**) Feature plots showing the expression of genes enriched in basal progenitor cluster. Arrowheads indicate expression of those markers within the mitotic cluster. (**H**) Cross-species comparison between mouse and shark developing telencephalic datasets using SAMap reveals a correlation trend across basal progenitors and apical progenitors of both species. (**I**) Lists of genes that show shared expression across species (up) and species specificity (down). Abbreviations: AP, apical progenitor; BP, basal progenitor; SVZ, subventricular zone; VZ, ventricular zone.

To further explore AP diversity, we isolated, reclustered and regressed the effect of the cell cycle (methods) of the AP clusters (including mitotic BPs in the analysis), revealing transcriptional variability along the pallial-subpallial axis. In the new embedding, the original M-phase clusters, redistributed through this axis within their respective cell type identities (fig. S4B). Based on known markers in other vertebrates and their relative position in the dorso-ventral axis, we propose the existence of at least five broad AP subtypes (fig.S4B,C): medial ganglionic eminence (MGE)/preoptic area precursors (shh-high), lateral ganglionic eminence (LGE) precursors (meis2-high), dorsal LGE/pallial-subpallial border precursors (prdm16-etv1-high), ventrolateral pallium precursors (pax6-lrp4-high), and dorsomedial pallium precursors (lrp4-lhx9-foxp2-rspo1-high). Additionally, we detected a cluster of emerging BP-like cells expressing st18 and ebf3 (fig.S4B).

Along the pallial lineage, we observed the sequential expression of pax6, eomes (previously known as tbr2), and tbr1 (Fig. 4D). In mice, these genes form a direct feedforward genetic cascade that regulates the sequential differentiation of pallial APs (Pax6 high) into BPs (Pax6 low and Eomes high) and finally into early (Tbr1) (*27*). Immunohistochemistry confirmed high Pax6 expression in the VZ and lower levels in the SVZ, where Eomes transcript is specifically expressed (Fig. 4E). These findings align with previous observations (*2*) and reveal strong conservation of this genetic cascade in driving the early differentiation of telencephalic excitatory neurons. This conservation accordingly dates back at least to the last gnathostome common ancestor and suggests the presence of an equivalent BP state in the developing pallium of sharks.

One central question is whether this BP state in sharks represents a distinct progenitor type or just a transitional state towards differentiation. To investigate this, we examined cells with high BP scores in our single-nucleus dataset. These cells were distributed across both early differentiating branches and within progenitor clusters (Fig. 1C). We noticed that the mitotic gene expression signature grouped cells together into a “M-phase cluster” (Fig.4F), and that a proportion of BPs fell into this mitotic cluster (arrowheads in Fig. 1C). Likewise, top enriched pallial BP genes were co-expressed in the same mitotic cluster area than the one predicted by cell score analysis, confirming that a proportion of BPs must be mitotically active (arrowheads in Fig. 4G). In summary, BP identity in our dataset likely represents a state encompassing cells which just exited the cell cycle from AP state and are now entering into BP state, as well as actively dividing BPs and cells transitioning from BP state towards differentiation. The presence of mitotic cells (PH3-positive) cells in a subventricular position, noted in our previous work (*2*), further supports the existence of a distinct BP progenitor state in sharks. Among the pallial BP-enriched genes, we identified the transcription factor Ebf3, which we validated by HCR to be highly expressed in the pallial svz, where Eomes is also expressed (Fig. 4E).

To unbiasedly assess potential homology between shark and murine BPs, we conducted transcriptional comparisons using published single-cell data (*13*, *24*) from the developing mouse telencephalon at equivalent embryonic stages (E12-14; Methods), where APs and BPs were easily identifiable. We annotated shark pallial mitotic BPs as a distinct cluster from mitotic APs, guided by BP marker expression profiles (Fig. 4G). Our cross-species comparison using SAMap revealed a clear correlation trend between shark and mouse APs and shark and mouse BPs (Fig. 4H).

Overall, these findings suggest that shark BPs closely resemble their mouse counterparts in transcriptomic profile, mitotic activity, and SVZ localization, supporting an evolutionary common origin of indirect neurogenesis dating back to early gnathostomes.

We next compared the transcriptomic profiles of APs and BPs across species. In order to capture robust transcriptomic signatures, we identified genes significantly enriched in each species (log2 fold change > 1.5). We then determined the set of shared and species-specific orthologous genes for each cell type. The shared gene signature revealed that APs in both species exhibit high expression of Notch-family genes, while BPs express the Delta gene Dll1. This suggests that the regulation of the transition from AP to BP state by the Notch-Delta pathway is a core ancestral feature.

Many of the commonly enriched orthologous genes between shark and mouse BPs corresponded with established BP markers in mammals. This conserved signature includes transcription factors such as Ngn1, Ngn2, Eomes, Insm1, and Lhx9 (Fig. 2I). Together, these findings suggest that the Delta-Notch signaling axis, combined with this core set of transcriptional regulators, constitutes an evolutionary toolkit for indirect neurogenesis that has persisted for over 450 million years. Additionally, shared genes among APs appear to regulate stem cell maintenance, Hedgehog/FGF signaling pathways, and chromatin remodeling (Fig. 2I).

Regarding species-specific signatures, mouse APs display strong specific expression of Sfrp2, a Wnt pathway modulator, which could reflect adaptations to the increased proliferative complexity required for layered cortical development. Mouse BPs exhibit an enhanced proliferative profile compared to their shark counterparts, characterized by the acquisition of genes such as Hes6 and E2f1. Additionally, the retention of Argap10 (a Rho GFPase involved in cytoskeletal organization and cell migration, commonly enriched in mouse and shark APs) in mouse BPs could be associated with the radial expansion necessary for neocortical layering. In contrast, shark BPs show a stronger differentiation profile, with elevated expression of Ebf3 and Wnt7b, suggesting more limited BP amplification relative to mice. Together, our results indicate that core developmental mechanisms underlying pallial indirect neurogenesis were already established in the common ancestor of all jawed vertebrates and point to molecular evolutionary changes that may have promoted the retention of proliferative capacity in mammalian BPs, thereby contributing to mammalian pallial expansion.

### Identification of Cajal-Retzius-like cells suggest their early emergence in vertebrate brain evolution

Building on our evidence of a conserved dorsal pallial sector and that mechanisms of pallial neurogenesis were probably already established in the last common ancestor of jawed vertebrates, we next asked whether additional, defining features of the telencephalon might also trace back to this deep evolutionary origin.

We focused on one small cluster of pallial neurons characterized by Tp73, Ebf3 and Lhx1 expression (Fig. 5A,B) which are the molecular signature of mammalian Cajal-Retzius cells (CRs) (*28*). CR cells form a heterogeneous population, predominantly of pallial origin, arising from discrete sources near the pallial– subpallial border before migrating to populate layer I of the developing cortex and hippocampus (*29*, *30*). They are a developmental transient cell type, which undergo apoptosis after cortical development is complete. CR cells are best known for producing reelin, a signalling molecule crucial for guiding the radial migration and proper lamination of neurons of the neocortex (*31*). The presence of CR homologous types in non-mammalian vertebrates is a matter of debate, with some evidence in reptiles, controversy in birds and no evidence to date in fish (*28*, *32–35*).

**Figure 5.**
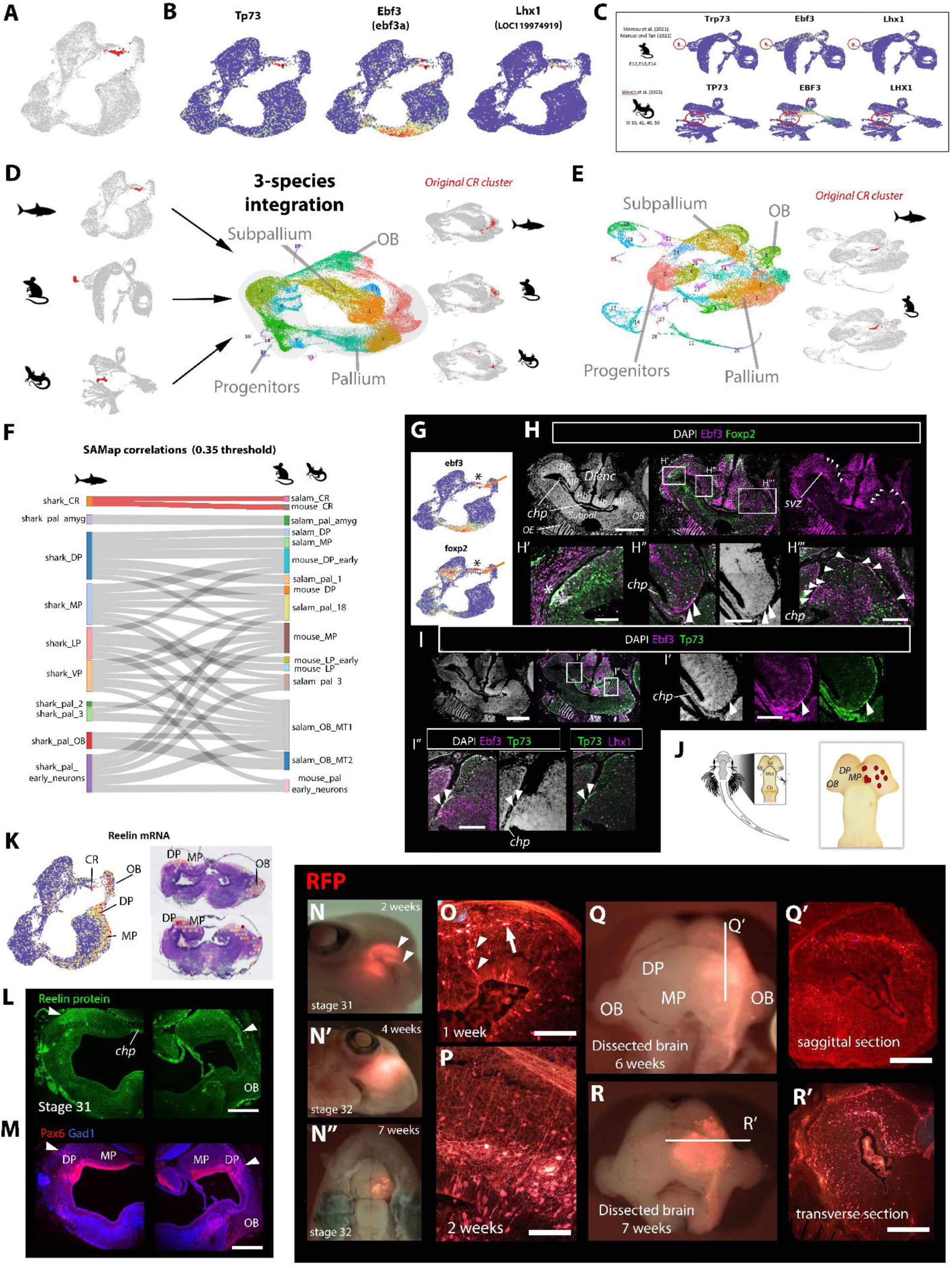
Characterization of a potential Cajal-Retzius (CR) homologous cell type in the shark embryonic telencephalon and identification of a layered structure in the pallium. (**A**) Umap of the whole shark dataset, highlighting in red the cluster of the putative CR cells. (**B**) Featureplot showing the expression of three genes enriched in the CR cluster, which are transcriptional signature of mammalian CRs. (**C**) Same three genes showed specific expression in the CRs cluster of mouse and salamander datasets. (**D-E**) Integration of three (D) and two (E) species dataset into one common shared space where genes from each species were converted to their respective orthogroups at the vertebrate taxonomic level. (**F**) Sankey plot showing correlation across pallial mature clusters from shark (left column) versus mouse and salamander (right column). Highlighted in red is the correlation between the CR clusters across species. (**G-J**) Localization of CR cells in transverse sections of the embryonic brain at stage 31. (G) Feature plots showing the expression of ebf3 and foxp2. Asterisk indicate the upper cluster where both markers colocalize and corresponds to the pallial amygdala. Orange arrow indicates the CR cluster, which expresses ebf3 but not foxp2. (H-H’’) HCR in a transverse section of the shark telencephalon showing expression of ebf3 and foxp2. Insets in H represent the area showed at higher magnification in H’-H’’. Arrowheads indicate the position of CR cells throughout the pial surface of the pallium. (I-I’’) HCR in a transverse section of the shark telencephalon showing expression of ebf3 and tp73. Insets indicate the position if the area showed at higher magnification in I’ and I’’. Arrowheads point to cells showing colocalization of the markers indicated. (J) (Left) Schematic drawing of a dorsal view of a shark embryo at around stage 30-31 and the position of the brain. (Right) Representation of a dorsal view of the anterior embryonic shark brain. Red circles represent the position of the identified CR-like cells. (**K**) Reelin transcript expression in the single nucleus (left) and spatial dataset (right). (**L-M**) Transverse section through the telencephalon of a stage 31 embryo showing the expression of Reelin (L) and Pax6-Gad1 (M) proteins. (**N-R’**) Injection and electroporation of a CMV-RFP reporter plasmid in the telencephalon of stage 29 embryos labels a neuronal population in the dorsomedial telencephalon organized in a layer. (N-N’) External view of the head of catshark embryos at different times after electroporation. Arrowheads show apparent streams of migrating neurons in the dorsal pallial area. (O-P) Sections through the pallium one week (O) and two weeks (P) after electroporation shows RFP-positive neurons emerging from the ventricular zone (arrowheads) and extending tangentially into an incipient layer. (Q-R’) 6-7 weeks after electroporation, strong RFP labelling appears in the dorsal telencephalon forming a noticeable neuronal layer visible in both sagittal (Q’) and transverse (R’) sections. Lines in Q and R indicate the approximated section level of Q’ and R’, respectively. Abbreviations: Cb, cerebellum; chp, choroid plexus; Dienc, diencephalon; DP, dorsal pallium; Hb, habenula; Mid, midbrain; MP, medial pallium; OB, olfactory bulb; OE, olfactory epithelium; SVZ, subventricular zone; Tel, telencephalon. Scale bars: 50 μm in (P,O); ; 200μm in (L,M, Q’,R’).

To test whether our potential CR cluster showed transcriptomic similarity beyond the three key markers across other species, we used the mouse integrated dataset in which a CR cluster was clearly defined in the original annotations (Fig. 5C)(*13*, *24*). After reprocessing the salamander dataset (methods)(*5*), we found a non-annotated cluster specifically expressing Tp73, Ebf3 and Lhx1 (Fig. 5C), which we annotated as potential CR salamander homolog and included in our three-way comparison. After integrating the three datasets based on shared orthogroups (methods, Fig. 5D, fig.S5B,C), we observed that CR-like cells from the three species co-distributed closely in the new embedding (Fig. 5D) and belonged to the same integrated cluster (fig. S5B). When integrating shark and mouse datasets, both CR types found each other in one of the most robust cluster of the whole dataset (Fig. 5E). We performed additional independent integration methods that yielded similar results. SAMap comparisons showed that the CR cluster was the amongst the few cell types that found a strong reciprocal correlative across the three species (red lines in Fig. 5F and red arrowheads, fig.S5A). Taken together, these results provide strong evidence that CR homolog cell types exist in sharks and salamanders, suggesting a far more ancient origin of CRs in vertebrates than previously hypothesized.

We next examined the localization of CR cells in the developing shark telencephalon. No conclusive CR cell localization was achieved with spatial transcriptomics using the three markers, likely due to the 50 μm resolution and the difficulty of detecting a rare, potentially anatomically dispersed cell type. To overcome these limitations, we performed HCR detection for the main CR markers on transverse sections of stage 30 shark telencephalon. To distinguish potential CR cells from another small, uncharacterized pallial population identified in our UMAP analysis, we selected Ebf3 and Foxp2 as differential markers. In the single-cell dataset, the putative CR cluster expressed Ebf3 but not Foxp2 (orange arrows in Fig. 5G), whereas the other population was Ebf3-Foxp2-double positive (asterisks Fig 5G). Using HCR, we found Ebf3+/ Foxp2+ cells located between the lateral and ventral pallium (asterisk Fig. 5H’). Cross-species comparison showed a strong correlation of the double-positive cluster cells with the salamander pallial amygdala cluster (green arrowheads, fig. 5SA), indicating a possible homolog in shark.

Ebf3-positive/Foxp2-negative cells were detected in the subventricular zone (svz in Fig.5H) likely representing basal progenitors rather than CR cells, consistent with results presented before showing Ebf3 expression in BPs (Fig.4E). The only other evident Ebf3+/Foxp2– cells occurred along the pial surface of the pallium, most prominently near the choroid plexus attachment in medial regions (arrow heads in Fig. 5H,H’’), showing analogous distribution to CR in mouse and human (*28*, *31*). At more caudal levels, they extended from medial pallial regions toward areas adjacent to the olfactory bulb (arrowheads in Fig. 5H’’’). We also examined the CR marker Lhx1. Only a few neurons in the caudal medial pallium expressed the three markers (Ebf3+/Tp73+/Lhx1+) (Fig. 5I’’), suggesting that CR subtypes may have distinct distributions across the telencephalon.

In summary, cells expressing CR markers were observed near the choroid plexus attachment in medial and caudal regions of the medial pallium, and extended along the pial surface of the dorsal and lateral pallium to areas adjacent to the olfactory bulb (Fig. 5J). Our results suggest that the molecular profile and anatomical distribution of CR-like cells are highly conserved from sharks to mammals.

We next investigated the expression pattern of Reelin, a secreted extracellular matrix protein known to be produced predominantly, though not exclusively, by CR cells in mammals (*28*, *31*) and essential for regulating neuronal migration and cortical layer formation (*36*). In the single-nucleus dataset, reelin transcripts were enriched at the tip of the CR cluster, as well as in the olfactory bulb and part of the dorsomedial pallial clusters (Fig. 5K). This spatial pattern was confirmed both in our spatial transcriptomic data and via fluorescent immunohistochemistry detecting Reelin protein (Fig. 5K, L). To delineate pallial regions and identify tangentially migrating gabaergic neurons, we used Pax6 and Gad1 protein distributions, respectively (arrowheads, Fig. 5M). Interestingly, Reelin localized to areas harbouring CR cells, including the marginal zone of the dorsolateral pallium and the olfactory bulb (Fig. 5K, L). Consistent with our previous findings (*1*), Reelin expression also overlapped with migratory interneurons entering the pallium through dorsal subpial regions (arrowheads, Fig. 5L,M). Given its role as an extracellular matrix molecule orchestrating neuronal positioning and migration during mammalian telencephalic development, Reelin’s conserved spatial association with CR cells and migratory neurons in shark supports evolutionary conservation of its developmental functions.

Intrigued by the developmental fate of the shark dorsal pallium, we electroporated a non-integrative expression plasmid (CAG-CMV-RFP; see Methods) into the telencephalon of five stage 29 embryos and monitored them over 4–7 weeks (Figs. 5N–R). The most mature reached late embryonic stage 32, when telencephalic cytoarchitecture begins to resemble the mature brain. In all cases, RFP-positive cells consistently labeled the dorsal and lateral pallium, with occasional labeling in small regions of the lateral subpallium (LGE). One-week post-electroporation, labeled cells were observed emerging from the ventricular zone with radial migratory morphology (arrowheads, Fig. 5O) and in the mantle zone oriented tangentially (arrow, Fig. 5O). By two weeks (stage 31), migratory cell streams were visible along the dorsal pallium in live embryos (arrowheads, Fig. 5N); sectioned tissue revealed numerous axons and somata with both radial and tangential orientations, the latter forming an apparent layer parallel to the ventricular surface (Fig. 5P). At 4–7 weeks (stage 32), labelled neurons organized into a layered structure in the medial-dorsal pallial area, evident in both sagittal (Fig. 5Q’) and transverse (Fig. 5R’) planes. This arrangement aligned with the position of CR-like cells, as apparent by comparing RFP labelling in Figs. 5N’’, Q, and R with the CR distribution in Fig. 5J. Consistent with this, we previously reported a potential layered distribution in

*S. canicula* dorsal pallial area, with Dlx2-positive gabaergic neurons located beneath a layer of Tbr1-positive glutamatergic neurons in late embryos (see Fig. 6B,C in (*1*)). Taken together, these observations reveal the emergence of a complex neuronal organization in the shark dorsal pallium by embryonic stage 31, spatially correlated with CR-like cells, gabaergic migratory neurons and Reelin expression, suggestive of a conserved scaffold for pallial lamination.

## Discussion

In this study, we present a single nucleus and spatial transcriptomics atlas of the developing shark telencephalon during key stages of neurogenesis and neural differentiation, capturing the developmental trajectories of pallial and subpallial cell types. This is an important resource for comparative developmental studies that will help to shed light on the origin and evolution of telencephalic cell types, in particular to identify developmental events that might have emerged during the transition from jawless to jawed vertebrates.

Our reconstruction of subpallial trajectories supports conservation of MGE and LGE but did not find any cell type that may correspond to a CGE, which may have evolved later as an extension of the MGE. Notably, the gabaergic neurons migrating to the shark pallium exhibit an LGE-like profile, in contrast to the MGE-derived interneurons in mammals, implying that MGE-derived pallial migration may be a later evolutionary innovation. This evidence is in line with previous research suggesting an evolutionary shift in cortical interneuron migration from the LGE to the MGE in mammals and possibly in birds (*37*). Whether the pallial-directed population we observe in sharks corresponds to a known basal vertebrate type or represents a lineage-specific feature remains to be determined. Our cross-species comparison of developing subpallial cell types in sharks and salamanders revealed that the shark BSA structure closely matches salamander striatal and amygdalar cell types, helping to resolve a long-standing debate and supporting the identification of this structure as homologous to the striatal portion of the basal ganglia and elements of the subpallial amygdala.

Notably, and based on their arrangement in the UMAP projection of our single-nucleus transcriptomic dataset, pallial cell populations appear organized along a continuum of cell identity rather than forming distinct, branching lineages, as is more evident in the subpallium. This pattern suggests that neuronal identity in the pallium may be determined or strongly influenced by signaling gradients. Importantly, we observed the same general trend in the single-nucleus datasets from mouse and salamander, supporting the idea that these parallel modes of neuronal specification—gradient-based in the pallium and lineage-based in the subpallium—may represent an evolutionarily conserved feature across vertebrates.

The analysis of developing pallial cell types in sharks reveals at least four molecularly defined and spatially segregated-although partially overlapping-broad populations. The spatial progression of the pallial cell type identities from medial to ventral positions, coupled with characteristic marker gene expression profiles, provides a molecular–anatomical framework for homologizing shark pallial territories with those of other vertebrates. Our cross-species transcriptomic comparisons with equivalent mouse and salamander datasets revealed broad conserved correspondences between the dorsal-medial pallial sectors of all three species and between their lateral-ventral sectors. Notably, the shark dorsal pallium shares significant molecular similarity with the dorsal and medial pallium of both mouse and salamander, including enrichment of transcription factors such as Cux2 and Pou3f2—canonical regulators of upper-layer cortical neurons in mammals. These observations suggest that a distinct dorsal pallium emerged early in gnathostome evolution. This raises the intriguing possibility that key developmental programs underlying dorsal pallial evolution were already in place more than 400 million years ago, providing a conserved scaffold upon which later vertebrate innovations—such as the expanded mammalian neocortex—were built.

The possibility of the existence of a distinct dorsal pallium in sharks prompted us to explore whether additional hallmark features of pallial development might also be present in this lineage. Our results indicate that this is indeed the case. First, the developmental sequence of excitatory neurogenesis in the shark pallium follows the highly conserved Pax6–Eomes–Tbr1 transcriptional cascade, a genetic program that in other vertebrates underlies the orderly transition from apical progenitors to basal progenitors and finally to post-mitotic neurons (*27*). The presence of basal progenitors in a discrete subventricular zone— transcriptomically and anatomically comparable to their mammalian counterparts—supports that the machinery for indirect neurogenesis was already established in chondrichthyans. Second, we find that shark pallium development includes a population of Cajal–Retzius (CR)-like cells with a conserved molecular signature, including Tp73, Lhx1, and Ebf3. We observed these cells from medial pallial regions and extending along the pial surface over medial and dorsal pallial sectors, reaching into the olfactory bulb— closely mirroring the distribution of Reelin expression. In mammals, CR cells orchestrate laminar organization through Reelin secretion, and we propose that their analogous positioning in sharks, particularly in the medial and dorsal pallium, suggests that the cellular and molecular scaffold for Reelin-mediated regulation of neuronal migration may have arisen well before the origin of amniotes. The evolutionary history of CR-like cells remains poorly resolved. To date, they have not been reported in fish, and their presence in birds (24) and other amniotes (*7*, *34*, *38*) remains elusive. Our comparative study provide evidence for their presence in salamanders, indicating that a homologous cell type might have already existed in the last common ancestor of all jawed vertebrates. This conclusion implies that CR cells represent a deeply conserved neuronal class—both in molecular profile and spatial organization—and that their absence in previous non-mammalian studies may reflect technical or sampling limitations. Systematic examinations across a broader range of vertebrate lineages will be essential to determine the full extent of their conservation, reveal potential lineage-specific modifications, and clarify the role these cells may have played in the early evolution of pallial architecture. Finally, the dorsal pallium of late-stage shark embryos shows an incipient layered arrangement of neurons in the same territory enriched for CR-like cells and Reelin expression. Although far simpler than the laminated cortex of mammals, this organization could reflect a basic template for layered architecture that may have been inherited by all jawed vertebrates.

Taken together the occurrence of a conserved neurogenic program, indirect neurogenesis via basal progenitors, Cajal–Retzius cells with Reelin domains, and an early form of lamination in sharks suggests that much of the developmental toolkit for building a complex pallium may have been already in place in the last common ancestor of gnathostomes. We hypothesize that the emergence of this integrated suite of features facilitated both the expansion and functional elaboration of the pallium in early jawed vertebrates, potentially providing a selective advantage during the predator-rich, increasingly competitive marine ecosystems of the Silurian–Devonian period approximately 450 million years ago.

Cartilaginous fish embryos, many of which are oviparous and amenable to experimental manipulation, represent a powerful emerging model for investigating the developmental origins of the vertebrate brain, offering access to early neurodevelopment while retaining key conserved features of vertebrate brain organization. Our identification of homologous pallial structures and cell types between sharks, mammals, and amphibians establishes a foundation for future investigations into the molecular and regulatory changes underlying pallial evolution. Unlike the nuclear or highly divergent pallial organizations seen in birds and teleosts, the shark pallium appears to develop via radial expansion, making cartilaginous fishes particularly well-suited for dissecting fundamental principles of pallial neurogenesis and evolution. With rapidly advancing tools in multi-omics, proteomics and functional genetics it will soon be possible to experimentally test developmental mechanisms and evolutionary hypotheses directly in this key phylogenetic lineage.

## Materials and Methods

### Animal use

Eggs of *S. canicula* were supplied by the Marine Biological Model Supply Service of the CNRS UPMC Roscoff Biological Station (France) or kindly donated by the Aquarium Finisterrae (A Coruna, Spain). Upon arrival, eggs were kept in sea water tanks under controlled conditions of 16°C, pH 7.5-8.5 and salinity of 35 g/L. Suitable measures were taken to minimize animal pain or discomfort.

Embryos were deeply anesthetized with 0.5% tricaine methane sulfonate (MS-222, Sigma, St. Louis, MO) in filtered sea water before proceeding with dissection or fixation. Embryos were staged based on external features following (*10*). For fixation, embryos were decapitated and immersed in 4% paraformaldehyde (PFA) in PBS, overnight at 4°C.

All animal experiments were carried out according to the guidelines of the Committee for Animal Welfare and Institutional Animal Care and Use (IACUC) under EMBL’s Policy on the Protection and Welfare of Animals Used for Scientific purposes.

### Sample collection and nuclei preparation

Deeply anaesthetized embryos were staged and immersed in cold sea water containing 0.5% of MS222. After telencephalic dissections, the tissue was flash-frozen in an Eppendorf tube immersed in liquid nitrogen and stored at -80°C. Single nuclei were isolated from fresh frozen tissue using a modified protocol from (*7*). Dissected tissue was placed in 1.5 ml Eppendorf LoBind tubes, and flash-frozen in liquid nitrogen. Frozen tissues were then placed on ice and homogenized in 300 µl of HB buffer (250 mM sucrose, 25 mM KCl, 5 mM MgCl₂, 10 mM Tris-HCl pH 8.0, 0.1% IGEPAL, 1 µM DTT, 0.4 U/µl Murine RNase Inhibitor [NEB], 0.2 U/µl Superase-In [ThermoFisher) using micro pestles (Axygen). Homogenization was performed on ice and completed within 5 minutes. Unlysed tissue fragments were pelleted by centrifugation at 100 × g for 1 minute at 4 °C. The supernatant containing nuclei was transferred to a new LoBind tube and centrifuged at 400 × g for 4 minutes at 4 °C. After discarding the supernatant, the pellet was washed and resuspended in 300 µl of fresh HB buffer and nuclei were pelleted again at 400 × g for 4 minutes at 4 °C. Finally, the nuclei pellet was resuspended in 50 to 100 µl of ice-cold Dulbecco’s PBS supplemented with 1 µM DTT, 0.4 U/µl Murine RNase Inhibitor (NEB), and 0.2 U/µl Superase-In (ThermoFisher). The suspension was passed through a 40 µm Flowmi filter to remove aggregates. Nuclei quality and concentration were assessed by staining with SYBR Safe DNA stain and imaging on a Zeiss Axio Imager using a C-Chip hemocytometer (Neubauer improved).

### Single Nuclei Capture and Library Preparation

Single nuclei capture and library preparation were performed using the Chromium Single Cell 3’ Gene Expression Kit (v3.1 chemistry) and the Chromium Controller (10x Genomics), following the manufacturer’s protocol with slight modifications. Approximately 15,000 nuclei were loaded per capture channel. cDNA amplification was performed with 15 cycles for samples at stage 29– 30 and 12 cycles for samples at stage 30–31, using a 3-minute extension time. Final libraries were quantified using a Qubit fluorometer, and fragment sizes were assessed using an Agilent Bioanalyzer. Libraries were sequenced with 250 million PE reads on an Illumina NextSeq 2000 using the following read configuration: 28 cycles for Read 1, 10 cycles each for the i5 and i7 indices, and 90 cycles for Read 2.

### Visium 10X Spatial Transcriptomics

One fresh dissected telencephalon at stage 31, was embedded in OCT mounting medium, cryosectioned into 14μm transverse sections and placed on a Visium Spatial Gene Expression slide from 10X Genomics. Samples were processed according to manufacturer’s instructions. Briefly, slides were fixed in methanol and stained with Hematoxylin and Eosin. Images were taken with a Nikon Ti2 at 20x magnification. The slide was processed using the Visium Spatial Gene Expression Reagent kit. Permeabilization time was tested and determined the optimal at 8 minutes. cDNA libraries were amplified using 12 cycles for samples at stage 30–31 with a 1-minute extension time. Final libraries were quantified using a Qubit fluorometer, and fragment sizes were assessed using an Agilent Bioanalyzer. Sequencing was performed on an Illumina NextSeq 2000 with the following read configuration: 28 cycles for Read 1, 10 cycles each for the i5 and i7 indices, and 90 cycles for Read 2 with 200 million PE reads per library.

Spatial transcriptomics data was processed using the Space Ranger (v2.0.1) software (10x Genomics) with default parameters. Raw sequencing data were aligned to the *S.canicula* reference genome, and spatially resolved gene expression matrices were generated. Quality control, normalization, and downstream analyses, including clustering and visualization of spatial gene expression patterns, were performed using Seurat (v5) in R. Default workflows and parameter settings were used for data filtering, dimensionality reduction, clustering, and spatial mapping.

Spatial transcriptomics data generated using the 10x Genomics Visium platform were processed and analyzed in R (v4.3.1) using the Seurat package (v5.1.0). Following standard quality control and normalization procedures, we kept two high quality sections. Spatially enriched genes were identified using the FindSpatiallyVariableFeatures() function with default parameters. This method detects genes exhibiting significant spatial variability across tissue sections by leveraging spatial autocorrelation statistics, thereby highlighting genes with non-random spatial expression patterns. The resulting list of spatially enriched genes was used for downstream analyses and interpretation of spatial gene expression architecture within the tissue.

CIBERSORTx (*39*) was used to digitally sort out relative cell type contributions within individual Visium spots using the single-nuclei data set as reference. To achieve this, a signature matrix was generated from the single-nuclei data set. Top 30 cell type markers were selected from the globally annotated data set (Pallium, Subpallium, Subpallial Progenitors, Olfactory Bulb, Choroid Plexus, MGE) and only unique markers among the resulting list were used to generate the signature matrix. Based on the signature matrix spots were sorted for these cell types which resulted in a matrix with relative abundance estimates (summing to 1 for each spot) of global cell types across each spot that were afterwards visualized as features on the spatial map (Fig. S1E). Before assessing the dominant identity to each spot, we applied a mild filtering where spots exhibiting p values higher than 0.4 were labelled as “unassigned”. Additionally, the cell type with the highest proportion within a spot needed to be 5% higher than the second hit and the matrix was scaled. Subsequently, the cell type with the highest proportion within each spot was assigned as spot identity and the distribution was smoothed based on nearest neighbors’ identities (CRAN package: ‘FNN’) across the spatial map to mark cell type regions (Fig. 1I). This overall strategy was applied for all Visium analyses in this study with only the signature matrix input being changed based on the cell type resolution of interest for pallial and subpallial regions.

### Genome alignment, scRNA-seq preprocessing and quality control

A mapping index was created with CellRanger v6.1.1 using the *S.canicula* v1.1 genome (GCF_902713615) with default parameters. Demultiplexed FASTQ files from snRNA-seq were mapped against the index with “cellranger count” using default parameters.

### Single nucleus RNA sequencing data processing

Data processing was performed using Seurat (*11*)(v5.1.0) in R (v4.3.1). Raw count matrices from three independent experiments were initially processed separately, with cells retained only if they contained between 200 and 6000 detected genes. Following quality control, individual Seurat objects were created and log-normalized. These objects were subsequently merged while preserving experiment-specific metadata identifiers. To address batch effects, we implemented a dual integration strategy: first applying SCTransform normalization with glmGAMPOI variance stabilization, followed by Harmony batch correction using experiment origin as the batch covariate across the first 50 principal components. The integrated dataset was clustered using the Louvain algorithm via the FindClusters() function, with subsequent dimensionality reduction and visualization using UMAP. Differential gene expression analysis between clusters was conducted through Wilcoxon rank-sum tests (minimum 25% cells expressing genes in either cluster, log-fold change threshold of 0.25, unless otherwise specified), followed by cell type annotation.

For the identification of pallial molecular signatures across the different pallial regions, we used a more stringent criteria to select true marker genes (log2FoldChange > 0.5 , follow by visual inspection retaining only those genes that showed high cluster specificity).

For the identification of transcriptomic profiles of APs and BPs, we retained only those exhibiting a log2 fold change equal or higher than 1.5, in order to capture robust conserved transcriptomic signatures across species.

### RNA velocity

Raw reads of the 10x sequencing libraries were mapped individually using STARsolo (v2.7.9)(*40*) with the soloFeature Velocyto following the recommendations for 10x libraries (-- soloMultiMappers Uniform --soloCBmatchWLtype 1MM_multi_Nbase_pseudocounts -- soloUMIfiltering MultiGeneUMI_CR --soloUMIdedup 1MM_CR --soloCellFilter EmptyDrops_C). Spliced and Unspliced data matrices obtained were combined across all samples and loaded into python to run scVelo (*41*) using 30 PCs and 30 neighbours for the moments’ calculation. The resulting velocity fields were projected onto the existing UMAP embedding to visualize the predicted future states and lineage relationships of individual nuclei. This analysis provided additional insight into differentiation trajectories and complemented pseudotime inference from Monocle3.

### Monocle trajectory inference

Single nucleus RNA-seq data were analyzed using Seurat (v5.1.0) and Monocle3 (v1.3.4) to reconstruct developmental trajectories. After preprocessing and clustering in Seurat, the dataset was converted to a Monocle3 cell_data_set object. Cluster identities, UMAP embeddings, and cell metadata from Seurat were transferred to Monocle3. Cells were clustered and a trajectory graph was learned using UMAP-based dimensionality reduction, with parameters adjusted to optimize graph structure. Cells were ordered in pseudotime, and differential gene expression along trajectories was identified using Monocle3’s graph_test function. Visualization of gene expression dynamics and pseudotime was performed with plot_cells and FeaturePlot.

### AP diversity study

Apical progenitor populations were subsetted based on cluster identities, and data normalization, scaling, and dimensionality reduction were performed using SCTransform and principal component analysis (PCA). Re-integration was performed using Harmony, followed by UMAP embedding, neighborhood graph construction, and clustering. Cell cycle effects were regressed out using ScaleData() with the vars.to.regress argument set to S and G2M scores. Marker genes for each cluster were identified using differential expression analysis. For trajectory inference, the processed Seurat object was converted to a Monocle3 cell_data_set using as.cell_data_set(). Cluster identities, UMAP embeddings, and metadata were transferred to the Monocle3 object, and cells were reclustered with cluster_cells() using UMAP reduction. Principal graph learning was performed with learn_graph(), with parameters such as ncenter, minimal_branch_len, prune_graph, and geodesic_distance_ratio adjusted for optimal trajectory structure. Cells were ordered in pseudotime using order_cells(), and pseudotime values were extracted with pseudotime(). Differentially expressed genes along pseudotime and at branch points were identified using graph_test(), and gene modules were detected with find_gene_modules(). Visualization of gene expression dynamics and trajectories was performed using plot_cells(), plot_genes_in_pseudotime(), and ggplot2-based plotting functions.

### Trajectory analysis along the subpallial lineage

Subpallial clusters were subsetted and re-clustered using the Louvain algorithm in Seurat (v5.1.0) to refine population heterogeneity. Trajectory inference was performed with Monocle 3 (v1.3.4) by converting the subsetted Seurat object into a CellDataSet using default parameters. A principal graph was learned over the precomputed UMAP space with learn_graph(), and pseudotime ordering was calculated by specifying a root node at the progenitor-enriched. Branch-specific differentially expressed genes were identified using graph_test() with Moran’s I statistic to prioritize spatially coherent transcriptional changes along inferred differentiation trajectories.

### Cell state and cell cycle scoring analysis

Cell cycle phase assignment was performed using the CellCycleScoring() function in Seurat (v5.1.0), which calculates S and G2/M phase scores for each cell based on the expression of canonical marker genes. For this analysis, lists of S and G2/M phase genes originally curated in mouse were translated to their shark orthologs using OrthoFinder (v2.5.4) and used as input. Cells were classified into G1, S, or G2/M phases according to their highest phase score, and these assignments were retained as metadata for downstream analyses. Similarly, cell state scores were computed using the AddModuleScore() function in Seurat, with gene sets representing four cell states (apical progenitor, basal progenitor, early neuron, late neuron) curated from a published mouse study (*13*) and translated to shark orthologs using OrthoFinder. The resulting module scores quantified the relative enrichment of each cell for the selected gene sets, facilitating the interpretation of cellular heterogeneity in terms of both cell cycle phase and developmental cell state.

### Cross-species comparisons using common orthogroups

We estimated the transcriptome of the last vertebrate common ancestor by assigning each modern gene to an orthogroup at the vertebrate taxonomic level with the emapper tool (v.2.1.3) (*42*) and convert each modern animal’s count matrix from a matrix of “modern gene” counts per cell, to a matrix of “orthogroup counts” per cell. In some cases, true lineage-specific gene duplications will therefore be assigned the same vertebrate orthogroup, and in such cases the counts for the true genes were summed to estimate the expression of the orthogroup. While this can obscure some of the differences between closely-related, recently-derived modern cell types that rely on novel gene duplications in the modern animal, it also enables us to focus on the similarities between species, whose common ancestor likely had a simpler set of cell types relative to either descendant lineage.

Proteomes for each animal were collected by and using each animal’s genome resources we kept only the longest protein encoded per gene, such that we had a single protein per gene. In the resulting simplified proteome, we then renamed each protein identifier to match the gene identifier that was used in the single cell RNAseq count matrix. We ran this simplified proteome through the emapper software (v.2.1.3) (*42*) with the command emapper.py -m mmseqs -i . For each unique peptide in the emapper output matrix, we obtained a gene identifier that matched the genes reported in the single cell count matrix of the relevant animal, and also an orthogroup assignment at a range of taxonomic levels, including for the vertebrate orthogroups relevant for this study. We used this information to relate genes to orthogroups for both species by creating a simple lookup table, with one column for genes and a second column with their orthogroup assignment.

### Cross-species comparison with SAMap

The single-cell objects were converted to the H5AD format using the “sceasy” R library (v0.0.7) and their metadata (gene and cell IDs) lightly edited to ensure cross-compatibility with SAMap. Pairwise all-against-all sequence alignments of the peptide sequences of the three species were conducted using MMseqs2 (commit 67949d7,(*43*)). Pairwise SAMap comparisons between shark and mouse as well as shark and salamander were performed (SAMap v1.0.15, (*20*)) using previously established consensus annotation for each species. The mapping tables of shark to mouse/salamander clusters were extracted from the corresponding SAMap object and concatenated with each other. The resulting matrix was visualized as simple and clustered heatmaps using seaborn v0.13.2 (*44*) and plotly v6.0.1, with cosine distance measures. For specific pallial and subpallial cell types comparisons across species, we generated heatmaps with the pheatmap R package, plotting SAMap correlation matrix values. Hierarchical clustering was applied to both rows and columns.

### Immunofluorescence

Three primary antibodies were used in this study. GAD (Glutamic acid decarboxylase) Polyclonal sheep anti-GAD65/67 provided by Dr. E. Mugnaini; Pax6 Cat. no. PRB-278P Polyclonal rabbit anti-Pax6 Covance, Emeryville, CA; RELN (Reelin; clone 142) Cat.no. MAB5366 Monoclonal mouse anti-RELN Millipore, Billerica, MA.

Briefly, a cocktail of primary antibodies at appropriate concentrations (GAD, 1:30000; Pax6 (1:200); Reelin (1:150) were incubated in PBS containing 0.5% Tween (PBST), overnight at room temperature. Secondary antibodies were incubated at a 1:100 concentration in a humid chamber for 2 hours at room temperature in the dark. We used Alexa 488-conjugated donkey anti-mouse; Alexa 546-conjugated donkey anti-rabbit, from Molecular probes Eugene, OR. Sections were rinsed in distilled water, allowed to dry and mounted in MOWIOL 4-88 Reagent (Calbiochem, MerkKGaA, Darmstadt, Germany).

### Hybridization Chain Reaction (HCR)

Cryosectioned tissue slides were thawed for 15 minutes at room temperature. Pre-hybridization was performed at 37°C for 30 minutes in a humidified chamber using pre-hybridization buffer supplemented with 10 mM VRC RNase inhibitor (5 μl per 100 μl buffer). Hybridization was carried out at 37°C with 8 nM probe concentration and 10 mM VRC RNase inhibitor in a humid chamber. Post-hybridization washes were performed at 37°C with decreasing ratios of wash buffer to 5xSSCT, followed by equilibration in 5xSSCT at room temperature. For signal amplification, hairpins were snap-cooled (2 μl of each 3 μM stock hairpin per 100 μl amplification buffer for a final 60 nM concentration), heated at 95°C for 90 seconds, and cooled to room temperature in the dark for 30 minutes. Amplification was performed by incubating slides overnight at room temperature in a humidified chamber with 100 μl of amplification buffer containing the hairpins. Slides were then washed in 5xSSCT, counterstained with DAPI (5 μg/ml final concentration, 10 minutes), washed, and mounted in Prolong Gold antifade (ThermoFisher scientific). Fluorescent images were taken in a Stellaris 8 confocal microscope.

### Plasmid injection an electroporation

Embryos at stages 29 and 30 were removed from their egg cases, deeply anaesthetized as indicated above and placed in cold filtered sea water containing 0.5% tricaine methane sulfonate (MS-222, Sigma, St. Louis, MO). We used a plasmid harbouring a CMV enhancer, Beta-acting promoter driving the expression of RFP. We injected 1 μl of plasmid dilution in the telencephalic ventricle (plasmid concentration of 200ng/μl) and electroporated into the telencephalic ventricular wall using flat electrodes flaking the head area. Electroporation settings (previously tested for maximizing efficiency and reducing damage) were as follows: 25V, pulse length 5 msec, interval 500 msec, 4 pulses, decay rate 10%, positive polarity (poring pulse); 10V, pulse length 50 msec, pulse interval 500 msec, 5 pulses, decay rate 40%, positive polarity (transfer pulse). We used a NepaGene electroporator (Nepa21 Type II).

After electroporation, embryos were immersed in oxygenated sea water and monitor their recovery. They were regularly photographed using a Leica M205 FCA (optics carrier – 12803510). Embryos were eventually deeply anaesthesized, sacrifized as indicated above, and processed for cryosectioning.

## Data/code availability

Code used for RNA mapping and SAMap comparison is available at http://git.embl.de/npapadop/scan-scrnaseq and archived at https://doi.org/10.5281/zenodo.15233987. The cross-species comparison data is available at https://doi.org/10.5281/zenodo.15234059.

Single nucleus and spatial transcriptomics data will be deposited to the Gene Expression Omnibus (GEO).

## Acknowledgments

The authors thank Kresimir Crnokic, Emily Savage and Meritxel Modejar Barba for their support in animal husbandry, Matthias Janeschik for valuable discussion and exchanges, Ahmad Al Alwash for invaluable help with the spatial transcriptomics protocol, Ram Reshef (University of Haifa) for providing the reporter plasmid, GeneCore and ALMF imaging facilities from EMBL. Sébastien Henry and Sophie Booker from Roscoff Biological Marine Station (CNRS, France), and the Aquarium Finisterrae (A Coruna, Spain) for providing catshark embryos.

## Author contributions

Conceptualization: I.Q-U, D.A.

Funding acquisition: D.A., I.Q-U

Planned experiments: I.Q-U, P.A.O, L.P., G.S., A.V., M.B., I.C.R-D, D.T-S.

Executed experiments: I. Q-U, P.A.O, L.P., G.S, A.V, M.B, D.T-S, I.C.R-D, L.S

Designed and executed plasmid electroporation experiments: I.Q-U, I.C.R-D, M.B, D.T-S

Data processing: P.A.O., T.G, N.P

Analysed data and gave critical input: L.S., N.P., T.G, H.K.

Supervision: D.A., H.K.

Interpreted the data and wrote the manuscript with input from all authors: I.Q-U,D.A.

## Data and materials availability

Single nucleus and spatial transcriptomics datasets will be deposited to the Gene Expression Omnibus (GEO).

