## Supplementary material for "An atlas of shark developing telencephalon reveals ancient origin of basal progenitors and Cajal-Retzius cells": Figures S1 to S5

**The PDF file includes:**

Figs. S1 to S5

**Other Supplementary Materials for this manuscript include the following:**

Data S1 to S5

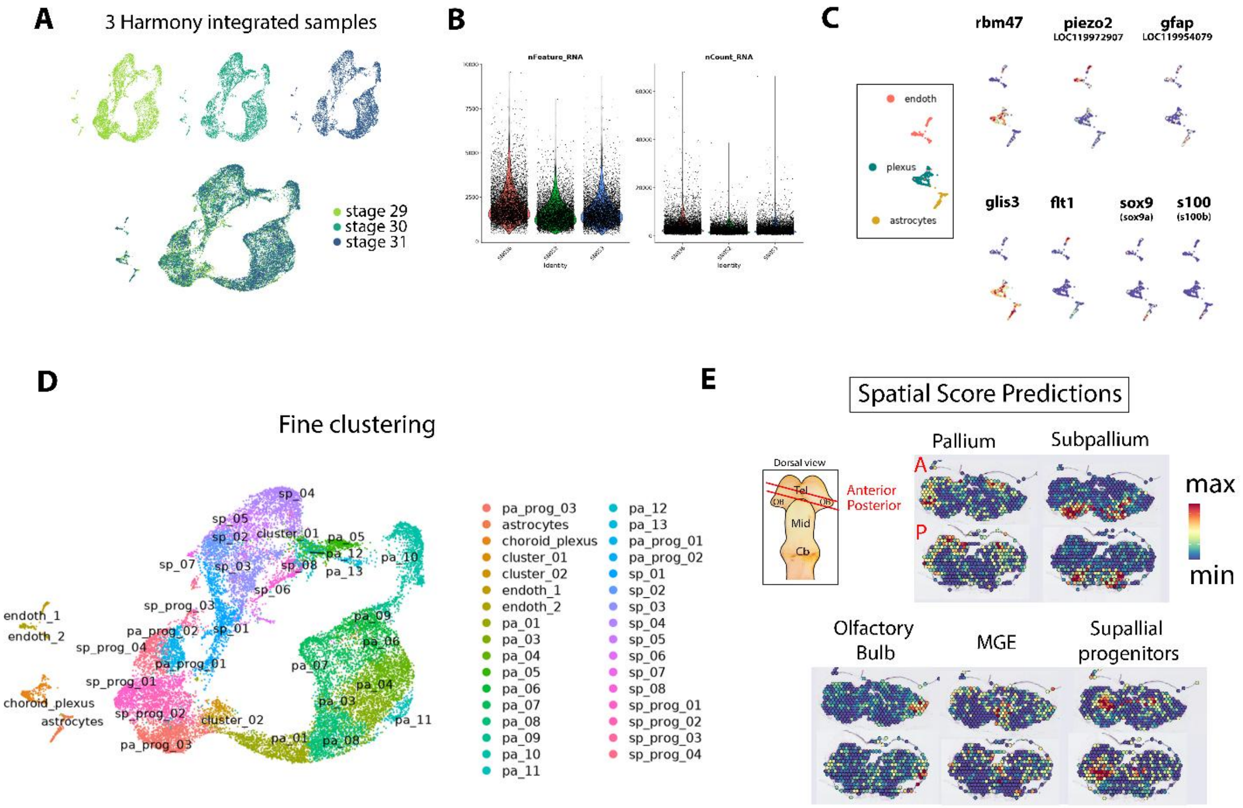


Fig. S1. **Quality control and exploration of the single nucleus and spatial transcriptomic dataset from the shark telencephalon.** (**A**) Umap showing the distribution of the three independent samples included in this study after harmony integration. (**B**) Violin plots showing features (left) and counts (right) distribution for each of the independent samples. (**C**) Identity of non-neuronal clusters revealed by selective marker expression. (**D**) Shark telencephalic dataset at a fine level of resolution. (**E**) Spatial score predictions for the main cell types in the telencephalon. Abbreviations: endoth, endothelial; MGE, medial ganglionic eminence; pa, pallium; prog, progenitors; sp, subpallium,

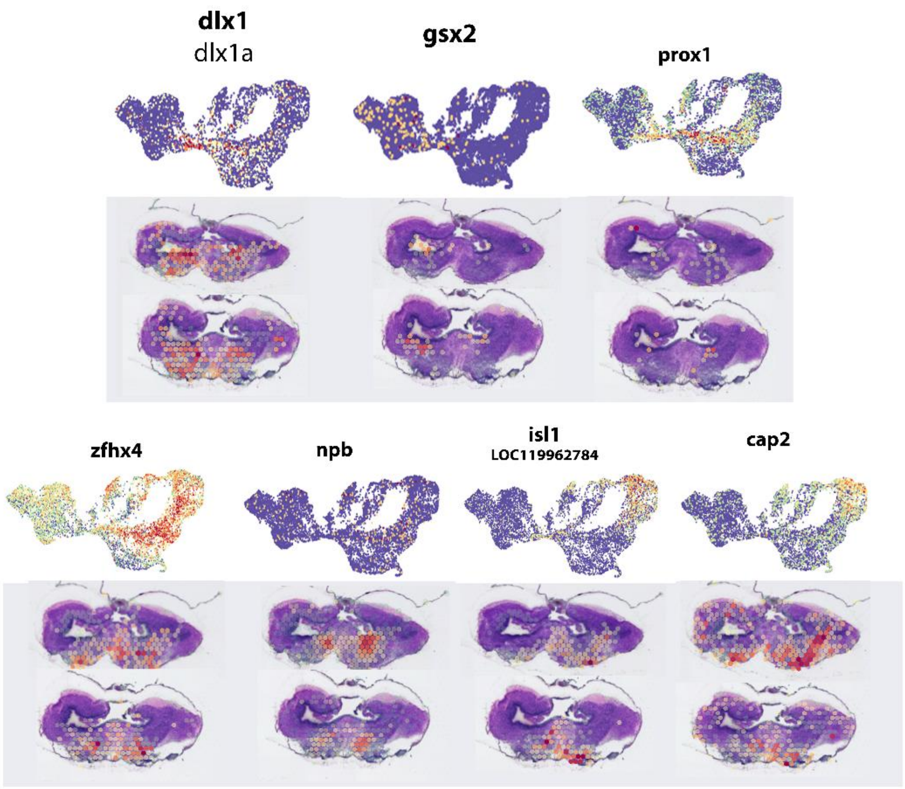


**Fig. S2. Expression pattern of selected subpallial genes in the subsetted subpallial branch (upper row) and in the Visium 10x spatial slides (lower row).**

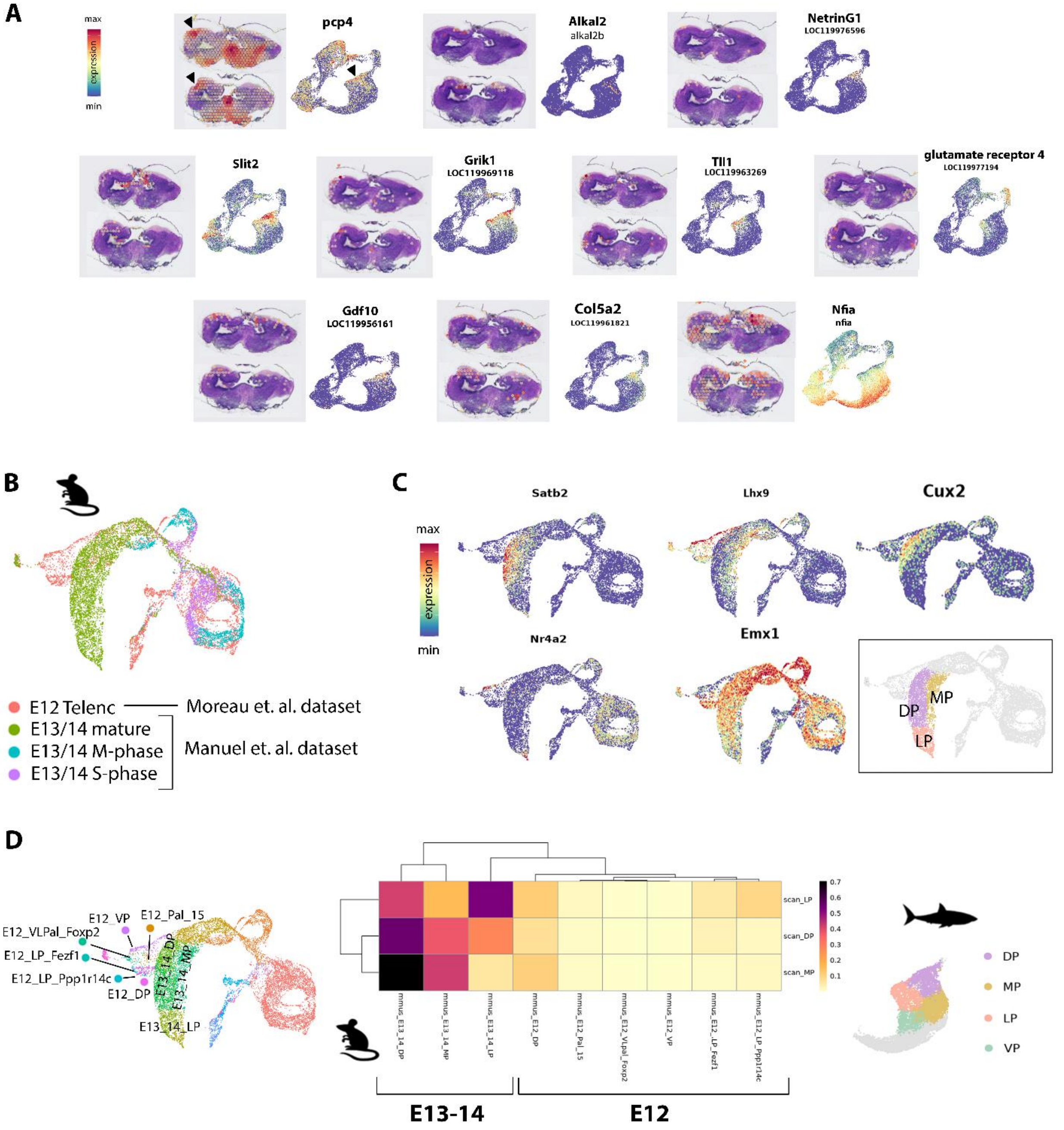


**Fig. S3. Study of the pallial cell types in the catshark telencephalon and cross-species comparison with mouse.** (**A**) Selected genes with specific expression in subregions of the pallium, shown both in the Umap and in the Visium 10x slides. (**B**) Integration of the two mouse datasets that were used to perform cross-species comparisons with the shark. (**C**) Annotation of the mouse pallial branch into their different subregions based on well-established marker genes. (**D**) Cross-species comparison of pallial clusters across shark and mouse shows the low correlation between the shark clusters and the mouse E12 maturing pallial neurons. Abbreviations: DP, dorsal pallium; LP, lateral pallium; MP, medial pallium; VP, ventral pallium.

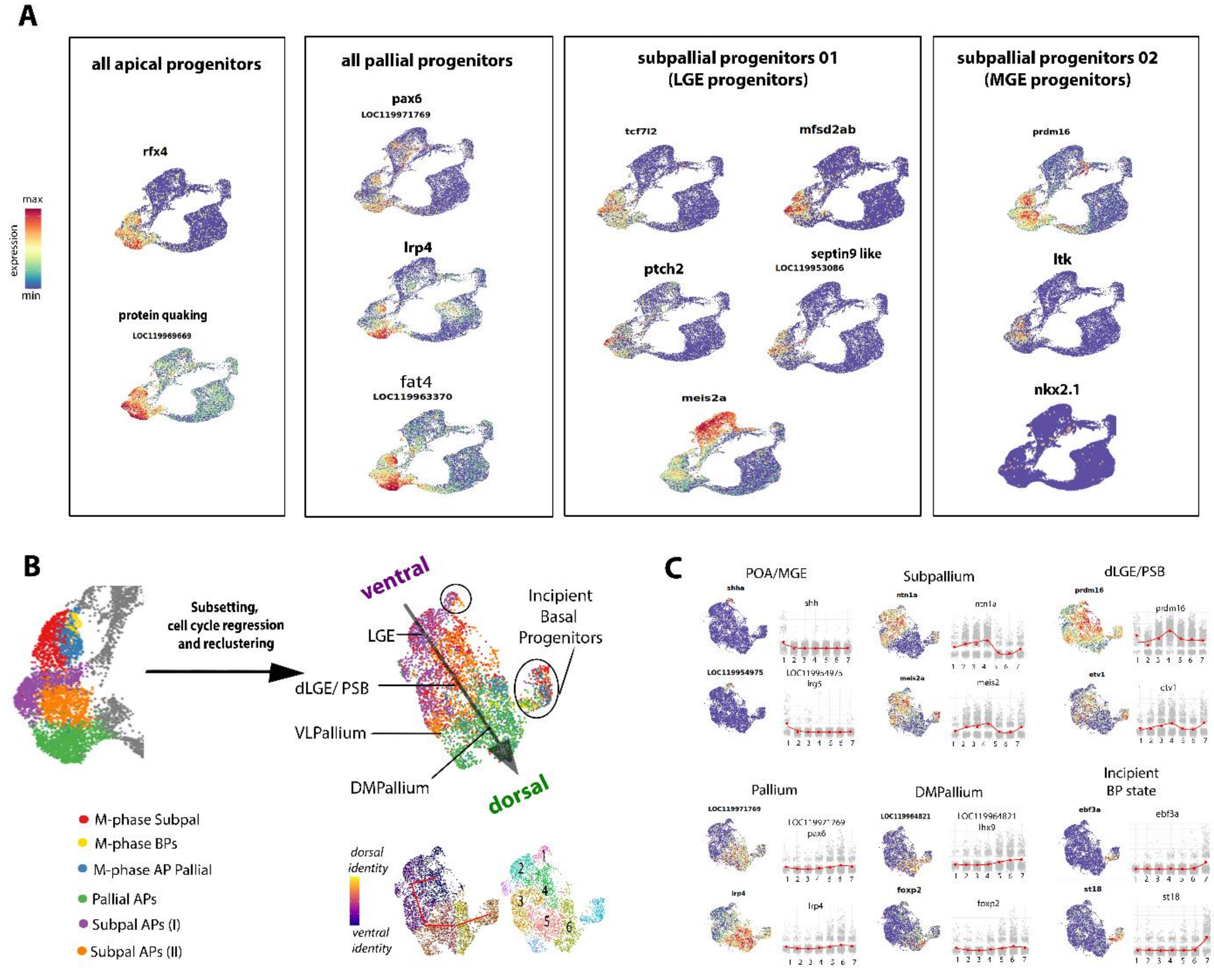


**Fig. S4. Study of diversity of apical progenitors in the catshark single nucleus dataset.** (**A**) Genes enriched on the different subsets of apical progenitors. (**B**) Study of apical progenitor variability after regressing the effect of the cell cycle. (Left) Cell cycle regression on progenitor clusters reveals variability along the dorso-ventral identity. (Right) (**C**) Expression of genes showing variability along the dorso-ventral axis. Abbreviations: BP, basal progenitor; Dlge, dorsal lateral ganglionic eminence; DMPallium, dorsalmedial pallium; MGE, medial ganglionic eminence; LGE, lateral ganglionic eminence; POA, preoptic area; VLPallium, ventral-lateral pallium.

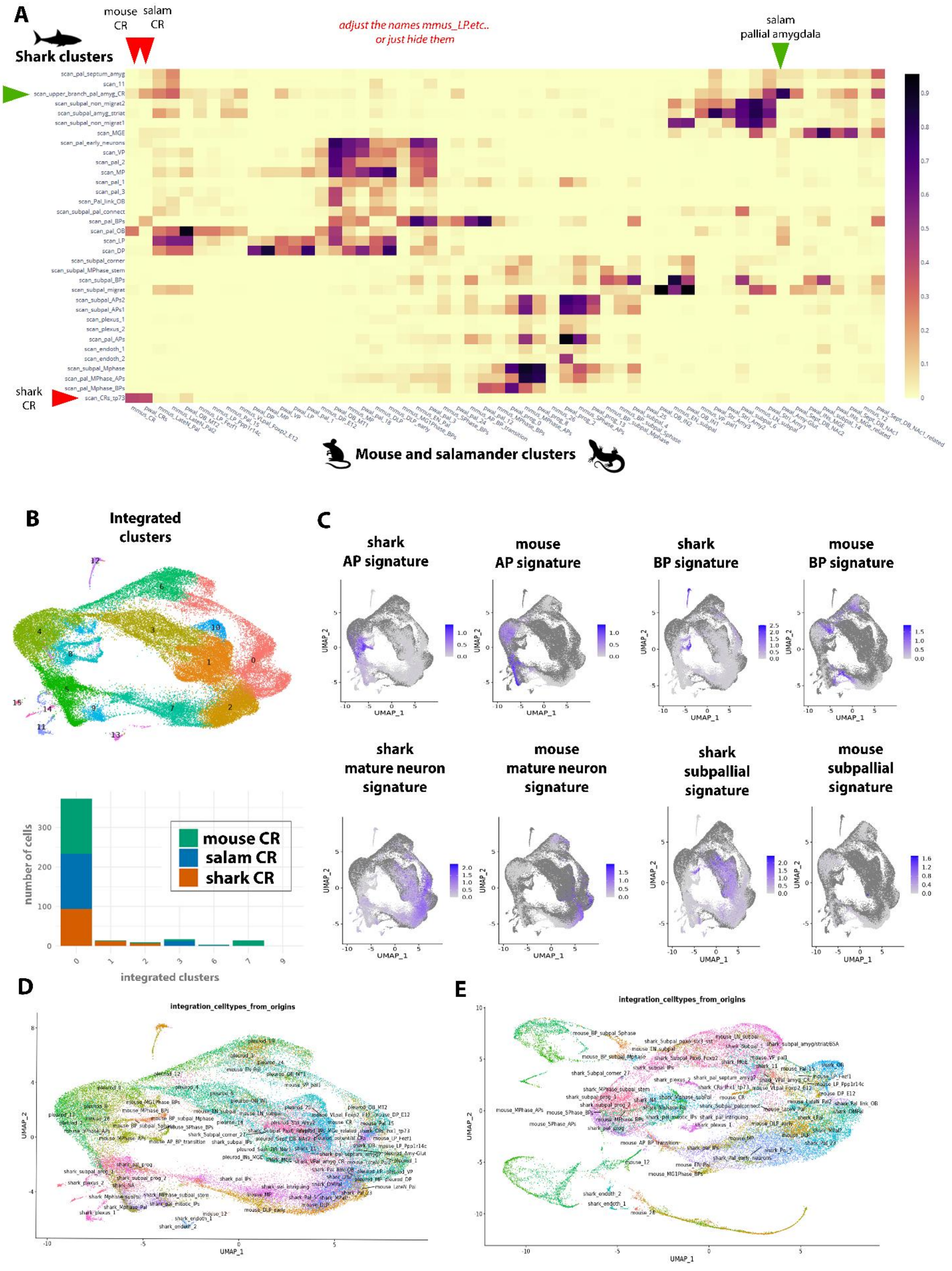
A
Shark clusters
mouse CR
salam CR
adjust the names mmus_LP.etc..
or just hide them
salam
pallial amygdala
shark CR
Mouse and salamander clusters
B
Integrated clusters
mouse CR
salam CR
shark CR
number of cells
integrated clusters
C
shark AP signature
mouse AP signature
shark BP signature
mouse BP signature
shark mature neuron signature
mouse mature neuron signature
shark subpallial signature
mouse subpallial signature
D
integration_celltypes_from_origins
E
integration_celltypes_from_origins
UMAP_1
UMAP_2

**Fig. S5. Cross-species comparisons of the whole datasets between shark, mouse and salamander.** (**A**) Heatmap sowing the correlation scores from SAMap analysis across the three species. Red arrowheads point to the CR clusters of the three species. Green arrowhead point to shark and salamander pallial amygdala clusters. (**B**) Three species integration using common orthogroups at a cluster resolution of 0.3 (top); distribution of cells belonging to CR clusters from the different species across the clusters in the 3-species integrated object. (**C**) Cell state signatures from shark and mouse projected on the three-species integrated object show the overall structure of the integrated object follows the individual object ones, with progenitors on the left, branching into pallial, subpallial and olfactory bulb clusters. (**D**) 3-species integrated object annotation showing original cluster names from each individual object. (**E**) 2-species (shark-mouse) integrated object showing original cluster names from each individual objects. Abbreviations: salam, salamander.

**Data S1. (separate file).** Differentially expressed genes across shark clusters. Coarse annotation.

**Data S2. (separate file).** Differentially expressed genes across shark clusters. Fine annotation

**Data S3. (separate file).** Differentially expressed genes across shark pallial areas

**Data S4. (separate file).** Differentially expressed genes across mouse pallial areas

**Data S5. (separate file).** Gene equivalences across shark and mouse. Orthofinder output
